# A method to remove the influence of fixative concentration on post-mortem T_2_ maps using a Kinetic Tensor model

**DOI:** 10.1101/2020.09.16.299784

**Authors:** Benjamin C. Tendler, Feng Qi, Sean Foxley, Menuka Pallebage-Gamarallage, Ricarda A.L. Menke, Olaf Ansorge, Samuel A. Hurley, Karla L. Miller

**Affiliations:** Wellcome Centre for Integrative Neuroimaging, FMRIB, Nuffield Department of Clinical Neurosciences, University of Oxford, Oxford, United Kingdom; Department of Radiology, University of Chicago, Chicago, IL, United States; Nuffield Department of Clinical Neurosciences, University of Oxford, Oxford, United Kingdom; Department of Radiology, University of Wisconsin - Madison, Madison, WI, United States

## Abstract

Formalin fixation has been shown to substantially reduce T_2_ estimates when performing post-mortem imaging, primarily driven by the presence of bulk fixative in tissue. Prior to scanning, post-mortem tissue samples are often placed into a fluid that has more favourable imaging properties, such as matched magnetic susceptibility. This study investigates whether there is any evidence for a change in T_2_ in regions close to the tissue surface in post-mortem T_2_ maps due to fixative outflux into this surrounding fluid. Furthermore, we investigate whether a simulated spatial map of fixative concentration can be used as a confound regressor to reduce T_2_ inhomogeneity. To achieve this, T_2_ maps and diffusion tensor estimates were obtained in 14 whole, formalin fixed post-mortem brains placed in fluorinert approximately 48 hours prior to scanning. This consisted of 7 brains fixed with 10% formalin and 7 brains fixed with 10% neutral buffered formalin (NBF). Fixative outflux was modelled using a Kinetic Tensor (KT) model, which incorporates voxelwise diffusion tensor estimates to account for diffusion anisotropy and tissue-specific diffusion coefficients. Brains fixed with 10% NBF revealed a spatial T_2_ pattern consistent with the modelled fixative outflux. Confound regression of fixative concentration reduced T_2_ inhomogeneity across both white and grey matter, with the greatest reduction attributed to the KT model vs simpler models of fixative outflux. No such effect was observed in brains fixed with 10% formalin. Correlations with ferritin and myelin proteolipid protein (PLP) histology lead to an increased similarity for the relationship between T_2_ and PLP for the two fixative types after KT correction. Only small correlations were identified between T_2_ and ferritin before and after KT correction.

## Introduction

Post-mortem imaging allows for the acquisition of high-resolution datasets and validation of the origin of image contrast through comparisons with histology. However, fresh tissue samples are vulnerable to damage through mechanical handling and decomposition though autolysis and putrefaction (1). To prevent this, samples are often first fixed prior to imaging using an aldehyde (2) solution such as formalin (3), to prevent decomposition and improve mechanical strength and stability.

Fixation has been shown to have a substantial effect on MR-relevant tissue properties, with decreases in T_1_ (4–9), T_2_ (4–11), T_2_* (6,7) and diffusivity (4,11–14) reported. These changes are thought to arise through either reactions with the aldehyde fixative solution in tissue via protein cross-linking (2) or presence of the bulk fixative solution (4) (fixative that has been absorbed into tissue). These changes have been shown to depend on fixative type, concentration, and vendor-specific composition (4,6,11).

To improve SNR, post-mortem samples are often first ‘washed’ via immersion in an external medium such as phosphate buffered saline (PBS), leading to exchange between the external medium and the bulk fixative solution. This process has been shown to restore T_2_ values close to those obtained prior to fixation (4), indicating that the change in T_2_ (due to fixation) is primarily driven by the presence of bulk fixative within tissue, rather than changes to the tissue itself. When considering formalin, the decrease in T_2_ has been estimated as linearly dependent on its concentration (4).

In addition to washing the post-mortem tissue samples, it has become increasingly commonplace to place tissue samples in an alternative fluid that has more favourable properties for imaging during scanning (15). One example is fluorinert (3M™), a susceptibility-matched perfluorocarbon fluid that produces no signal in MR images. This makes it possible to perform scanning without having to consider the signal from the surrounding medium (e.g. it is possible to perform imaging experiments considering a field-of-view that only covers the tissue sample) and obtain images that have minimal susceptibility-induced distortions or other artefacts.

Large samples (such as whole human post-mortem brains) are often not washed (16), due to the prohibitive length of time required for the external medium to penetrate into deep tissue. The time scales required for this can be inferred from literature examining brains undergoing immersion fixation with formalin (which has a similar molecular size), reporting multiple weeks for hemispheres/whole brains to become fully fixed (10,17).

Large samples can still be placed within an alternative fluid prior to scanning to improve the imaging environment (15). When considering formalin-fixed tissue, if there is any outflux of formalin into this surrounding medium, this may lead to a reduced concentration and a change in T_2_ in regions with close proximity to the brain surface. In this study, we investigate whether there is evidence for such an effect in T_2_ maps acquired in whole, formalin-fixed, human post-mortem brains placed in fluorinert approximately 48 hours prior to scanning. To achieve this, we simulate the outflux of fixative at the tissue surface and compare the resulting concentration distribution to the T_2_ values across our brain. We simulate outflux using a model that incorporates the effects of diffusion anisotropy and tissue specific diffusion coefficients, which aims to provide realistic modelling of fixative dynamics within different tissue types.

Previous studies of fixative dynamics (10,17) have aimed to characterise how the process of fixation and presence of fixative influences MRI parameters. Here, we take this approach one step further and propose that the resulting map of fixative concentration can be used as a confound regressor to account (or correct) for the effects of fixative concentration on T_2_. The correction is performed using a single global regressor that is fit to the T_2_ map across all of white matter. The importance of such a correction is evaluated by comparing the homogeneity of T_2_ estimates over white and grey matter separately within the post-mortem brains before and after correction. We evaluate this correction in a cohort of brains fixed with two types of fixative, 10% formalin and 10% neutral buffered formalin (NBF). Finally, resulting T_2_ estimates are correlated with histological measurements of ferritin and myelin proteolipid protein (PLP) content obtained within the same brain before and after correction.

## Theory

### The Kinetic Tensor model

Previous groups (10,17) have investigated the process of fixation by comparing MR estimates in tissue undergoing immersion fixation (influx of fixative) with mathematical models of fixative dynamics. For example, Yong-Hing et al. (17) modelled the influx of formalin fixative into a whole, human brain sample undergoing fixation by approximating the brain as a solid sphere and assuming a uniform isotropic diffusion coefficient. Dawe et al. (10) extended this approach by incorporating the geometry of the brain surface in hemispheres undergoing fixation. Here we build on this previous work by incorporating voxelwise diffusion tensor estimates (derived from diffusion MRI data from the same tissue sample) into our simulations (Fig. 1), known as the ‘Kinetic Tensor’ (KT) model. The KT model assumes that the concentration-driven diffusion of fixative can be modelled based on the self-diffusion process measured with diffusion MRI. We hypothesise that this will allow for more accurate modelling within different tissue types (e.g. grey and white matter) and incorporation of the orientation dependence of diffusivity estimates (due to diffusion anisotropy).

**Figure 1:**
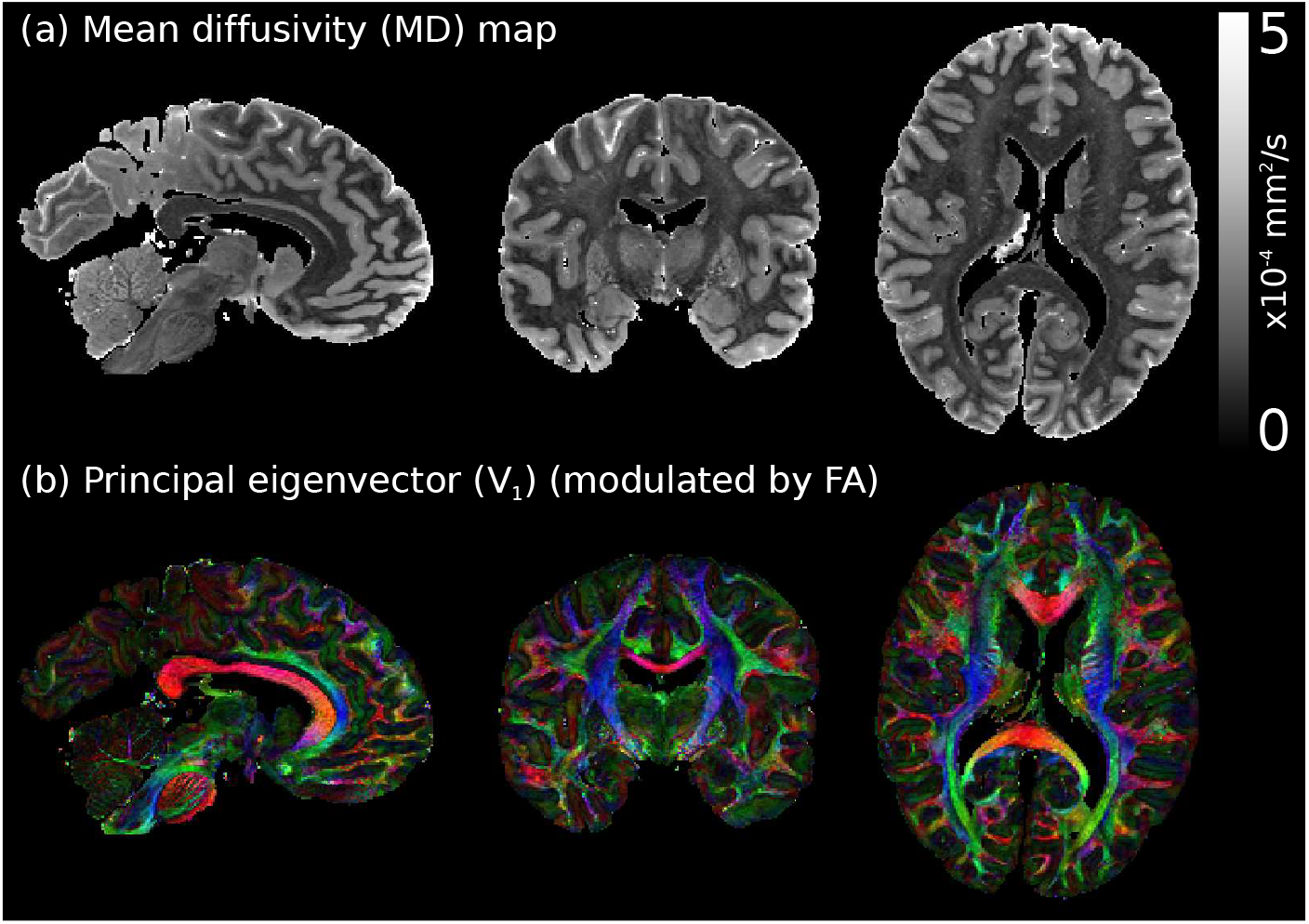
Diffusion tensor estimates in a whole post-mortem brain. Example diffusion tensor estimates from a single postmortem brain used in this study, displaying the (a) mean diffusivity (MD) and (b) principal eigenvector, 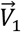, maps. Both grey and white matter have distinctive diffusivity estimates (a), and diffusion is highly anisotropic across the brain (b). The KT model incorporates these properties when modelling fixative dynamics. 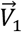 maps modulated by the fractional anisotropy (FA), where red: left-right, green: anterior-posterior, blue: superior-inferior. Diffusion imaging and processing protocol for this postmortem dataset is described in (18).

To achieve this, the concentration of fixative within tissue is simulated using Fick’s second law (19,20):

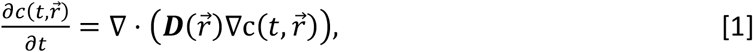

where 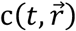 is the concentration of fixative at time *t* and position 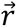, and 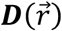 is the diffusion tensor:

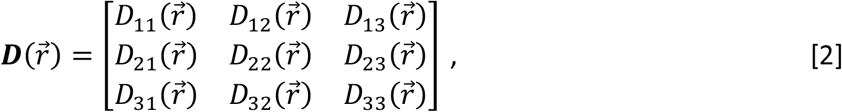

and the diffusion tensor is assumed to be symmetric (i.e. 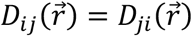). Given a set of initial conditions of the concentration distribution at *t* = 0, Eq. [1] can be evaluated. For example, the process of fixation (influx of fixative) can be modelled by assuming 0% fixative concentration within tissue and 100% fixative within the surrounding medium:

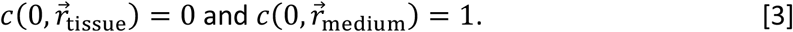

For the outflux of fixative (from tissue into the surrounding medium):

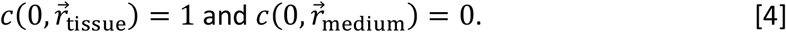

Here, we define *c* between 0 and 1, a unitless fractional concentration of fixative relative to the full concentration of the fixative solution.

### Incorporating realistic tissue geometries

Analytical solutions to Eq. [1] are only available when assuming simplified tissue geometries (e.g. approximating the brain as a sphere (17)). To incorporate realistic tissue geometries of the brain, Eq. [1] must be evaluated using an alternative means. Here we utilise a finite differences approach (as previously described in (19)) to model fixative dynamics within the brain. With finite differences, the spatial distribution of fixative is updated iteratively over a series of *n* time steps.

To achieve this, Eq. [1] is discretized and rearranged to solve for concentration 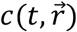 (Appendix Eq. [A1]). The spatial distribution of fixative concentration is subsequently simulated over a series of n time steps given a set of initial conditions (e.g. Eqs. [3] and [4]). Each time step estimates the change in concentration over the time period *τ* = *T/n*, where *T* is the total duration of the simulation. Figure 2 illustrates the simulated dynamics of fixative influx (initial conditions defined by Eq. [3]) into a whole post-mortem brain using the finite difference approach and the KT model. Fixative initially penetrates into the brain tissue through surfaces in contact with the fixative solution. Over time fixative moves further into the tissue, eventually leading to c = 1 across the entire brain.

**Figure 2:**
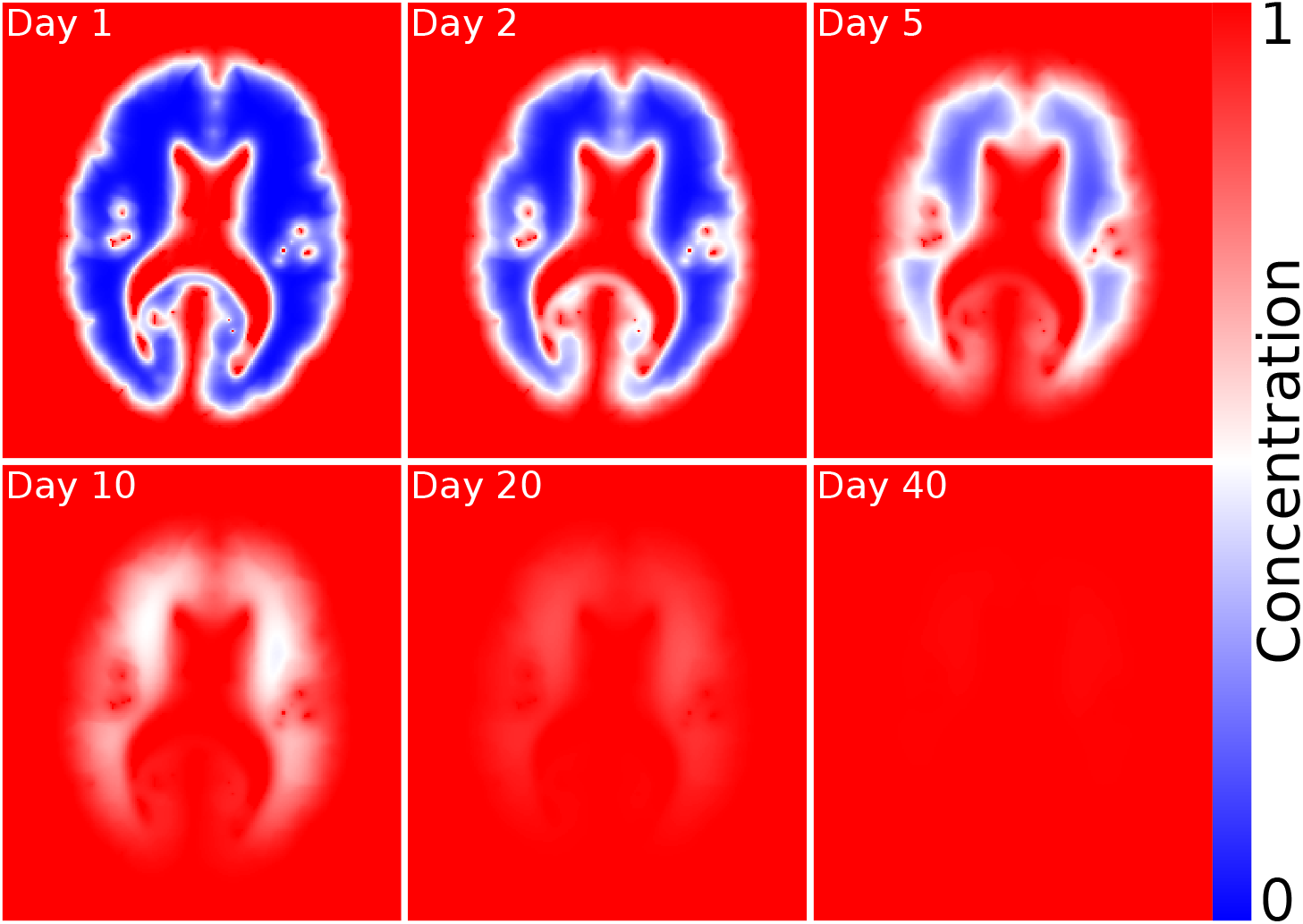
Modelling the influx of fixative using the KT model. Defining initial conditions from Eq. [3] (0% fixative concentration in tissue surrounded by 100% fixative), the KT model simulates the influx of fixative into tissue, accounting for both the relative diffusion coefficients of different tissue types and diffusion anisotropy (Fig. 1). Over time, fixative penetrates further into the brain, eventually leading to fully fixed tissue. For this brain sample, all voxels had a fixative concentration > 0.99 after 46 days, in broad agreement with a previous experimental observation reporting formalin fixation within ~38 days in a whole, human brain (17). Simulation performed using the diffusion tensor estimates in Fig. 1. Concentrations defined between 0 and 1, where 1 corresponds to a voxel containing 100% fixative.

### Kinetic tensor based confound regression

Prior to scanning, brain samples are typically removed from fixative and transferred into an alternative fluid that has more favourable imaging properties. This will lead to a concentration boundary at the brain surface (initial conditions defined by Eq. [4]), which may drive the outflux of fixative into the surrounding medium. Any outflux of fixative will lead to a decrease in its concentration in tissue and therefore a change in T_2_. When considering formalin, a previous study has estimated a 10-15 ms linear decrease in T_2_ per 2% concentration (4).

Figure 3 simulates the reduction in fixative concentration after modelling outflux (initial conditions defined by Eq. [4]) in a whole, post-mortem brain using the KT model. Initially, a reduced concentration of fixative is predicted within brain regions in close proximity to the brain surface, eventually leading to complete removal of bulk fixative within the brain after about 40 days. Large changes in concentration are observed near the brain surface within the first two days of immersion.

**Figure 3:**
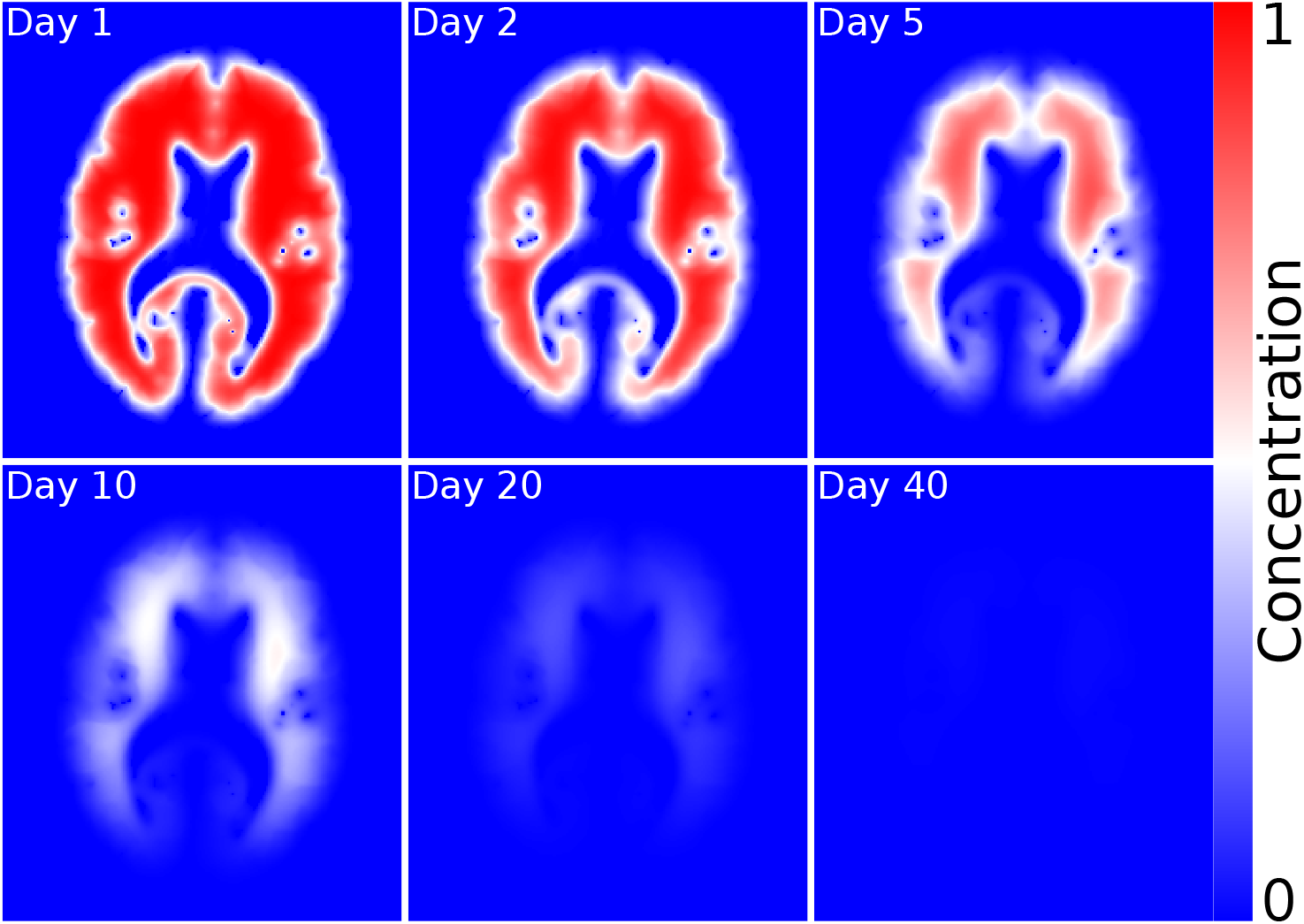
Modelling the outflux of fixative using the KT model. Defining initial conditions from Eq. [4] (100% fixative concentration in tissue surrounded by 0% fixative), here the KT model simulates the outflux of fixative into the surrounding medium. Over time the fixative concentration reduces throughout the brain, eventually leading to tissue with no bulk fixative solution remaining. For this brain sample, all voxels had a fixative concentration < 0.01 after 46 days. Simulation performed using the diffusion tensor estimates in Fig. 1. Concentrations defined between 0 and 1, where 1 corresponds to a voxel containing 100% fixative.

We propose using the resulting fixative concentration map derived from simulation as a confound regressor to account for the effect of fixative concentration on the quantitative T_2_ map. We perform this correction with the assumption that T_2_ varies linearly with bulk fixative concentration (4) (Fig. 4), defining:

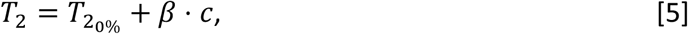

where *T*_2_0%__ is the T_2_ estimated at 0% bulk fixative concentration and *β* describes the rate of change of T_2_ with bulk fixative concentration. Here we perform this correction as a global regression, estimating a single *T*_2_0%__ and *β* per brain. From the estimate of *β*, we can subsequently perform a voxelwise regression of the bulk fixative concentration to generate a *T*_2_0%__ map; that is, the predicted T_2_ map in the absence of bulk fixative.

**Figure 4:**
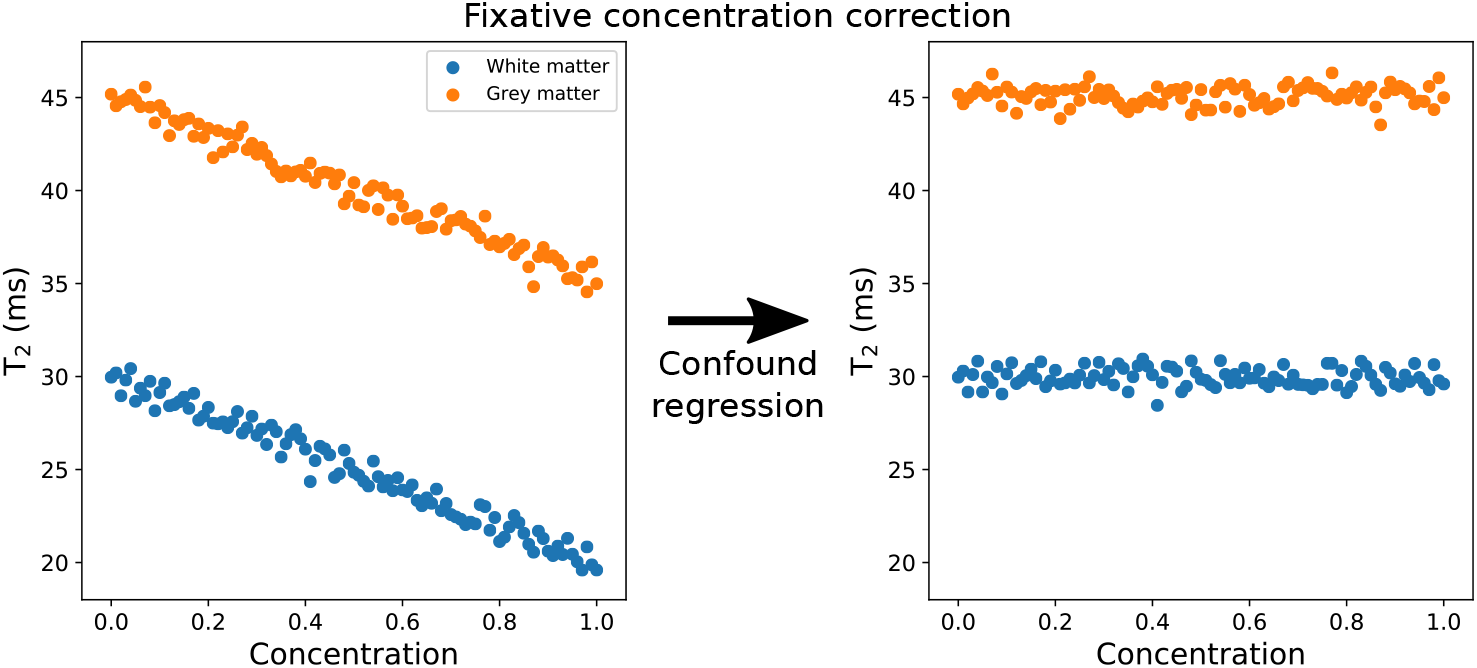
Simulation of the proposed confound regression of fixative concentration on T_2_. By simulating the outflux of fixative within tissue (e.g. Fig 3), we can plot the relationship between T_2_ and fixative concentration from the resulting concentration map (left). Here we assume that this will take the form of a linear relationship, as previously noted in (4). We can subsequently estimate and regress out the fixative concentration from the T_2_ estimates using Eq. [5] (right). Here we choose to perform this regression to estimate T_2_ at 0% bulk fixative concentration (*T*_2_0%__).

We compare this *“kinetic tensor”* (KT) correction to similar global regressions based on two other models: (i) a *“kinetic isotropy”* (KI) correction that assumes isotropic diffusivities, and (ii) a *“distance-to-surface”* (D2S) correction that considers only how close each voxel is to the nearest surface. The D2S model is a phenomenological correction that does not model fixative per se, but captures a simple geometric feature that relates to the flux of fixative.

## Methods

### Data acquisition and processing

Fourteen whole, formalin-fixed, postmortem brains (consisting of 11 brains from patients diagnosed with amyotrophic lateral sclerosis and 3 controls) were used in our experiment. Postmortem brains were extracted from the skull and immersion fixed in formalin (mean post-mortem delay = 3 ± 1 days, minimum = 1 day, maximum = 7 days). All brains were formalin-fixed for at least one month (mean duration = 125 ± 60 days, minimum = 45 days, maximum = 283 days) prior to scanning. Of these 14 brains, 7 were fixed in 10% formalin and 7 were fixed in 10% neutral-buffered formalin (NBF). The 10% formalin solution was made in-house by diluting 40% formaldehyde (Genta Medical, UK) in water (neutralised using marble chips). The 10% NBF solution (Genta Medical, UK) consisted of formaldehyde diluted in phosphate buffered saline (PBS). Details of individual brains are provided in Supporting Information Table S1.

Prior to scanning, excess formalin was removed from the brain surface and drained from the ventricles. Brains were subsequently submerged in fluorinert FC-3283 (3M™), a susceptibility matched fluorocarbon-based fluid that generates no MR signal, used to improve imaging quality. After removal of air bubbles/filling of the ventricles with fluorinert, brains were placed inside a custom scanning container filled with fluorinert. For full details of the brain packing process, see (21). All brains were immersed in fluorinert for approximately 48 hours prior to scanning.

Brains were scanned using a multi-echo turbo spin-echo (TSE) sequence on a 7T whole body Siemens system (6 echoes, TE = 13, 25, 38, 50, 63, 76 ms, TR = 1000 ms, resolution = 0.9 x 0.9 x 0.9 mm^3^, bandwidth = 166 Hz/pixel, turbo factor = 6), where each TE was obtained in a separate acquisition (time per acquisition = 36 minutes). These represent the typical imaging parameters for our T_2_ imaging protocol; the exact parameters evolved over the time-course of our experiment. Full details of the parameters for each sample are provided in Supporting Information Table S2. To account for any small changes in brain position between TEs, coregistration was performed using FSL FLIRT (22) (6 degrees of freedom transformation), though this typically led to no observable change in the resulting images.

When performing T_2_ mapping, B_1_ inhomogeneity can cause the signal to deviate from mono-exponential decay due to incomplete refocusing of echoes. As our 7T data was observed to demonstrate this effect, quantitative T_2_ maps were derived through voxelwise fitting of the signal using an extended phase graph (EPG) model that includes estimates of the B_1_ profile (23–25). This fitting is based on EPG software associated with (23) (available via contacting Matthias Weigel at). Full details of our EPG T_2_ fitting implementation is provided in Supporting Information.

While one might base KT modelling on a diffusion tensor atlas, in this case we have access to diffusion MRI for each individual brain sample being studied. Diffusion MRI data were acquired in each post-mortem brain using a diffusion-weighted steady-state free precession (DW-SSFP) sequence (26–30), from which diffusion-tensor estimates (three eigenvectors, 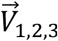, and three eigenvalues, *L*_1,2,3_) were derived over the whole brain at an effective b-value (b_eff_) of 4000 s/mm^2^. Details of the full acquisition protocol and processing pipeline for the diffusion data to a single b_eff_ are described in (18), and the full post-mortem protocol is described in (31). Example diffusion tensor estimates for a single post-mortem brain used in this study are displayed in Fig. 1.

### Modelling the outflux of fixative

The concentration of fixative within each brain was simulated assuming outflux into the surrounding medium for 48 hours (2000 time steps, *τ* = 86.4 s) using Eq. [A1], with initial conditions as defined in Eq. [4]. Throughout the simulation the concentration of fixative in the surrounding medium was kept constant 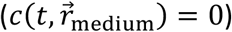. Although experimentally outflux will lead to a small concentration of fixative in the surrounding medium, given the time frame of our experiment (48 hours) and the volume of the surrounding medium used in our experiments, we believe this is a reasonable assumption. To prevent artefacts in the resulting simulations, voxels with spuriously high diffusion coefficients (empirically determined as > 1 · 10^−3^ mm^2^/s), were set equal to the mean of the surrounding tissue.

For the KT model, voxelwise diffusion tensors 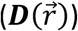 estimated over each postmortem brain (Fig. 1) were fed into Eq. [A1]. To assess the importance of incorporating diffusion anisotropy and voxelwise diffusion coefficients, two alternative models were investigated:

1. The KI model, which assumes isotropic uniform diffusion throughout tissue. To perform this, the diffusion tensor 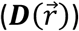 in Eq. [2] was substituted with a diagonal matrix, with each diagonal component set to the average mean diffusivity over the entire post-mortem brain.
2. The D2S model, which assumes the concentration of fixative in any given voxel is proportional to its distance (in mm) to the nearest surface. This model assumes a simple linear relationship between fixative concentration and the distance to surface (as opposed to accounting for fixative dynamics within tissue). The D2S model was calculated using the *distancemap* function in FSL (32,33).

Code for the KI and KT model used in this study is available at https://github.com/BenjaminTendler/KT_model.

### Fixative correction

The simulated fixative concentration maps were removed as a confound from our T_2_ maps by fitting with Eq. [5]. Fitting was performed as a single global regression, estimating a single value of *T*_2_0%__ and *β* per brain. One concern in fitting is potential tissue-type bias. Grey and white matter tissue are characterised by different T_2_ values and exhibit a spatial pattern that varies from centre to periphery (Supporting Information Fig. S4). Hence, it is likely that the true T_2_ maps will to some degree correlate with fixative models, since they share this general spatial distribution. To eliminate tissue-type bias on our fitting, *T*_2_0%__ and *β* were estimated from white matter voxels only. White matter masks were generated using FSL FAST (34) from the L3 diffusion tensor estimates. Both the concentration maps and tissue masks were estimated in the diffusion space of the postmortem brains, and transformed to the space of the T_2_ maps using FSL FLIRT (22) (6 degrees of freedom, estimated from the unprocessed TSE and DW-SSFP b0 data). It’s worth noting that as post-mortem experiments are not restricted by the same time constraints typically encountered in-vivo, diffusion scans can be performed with a low bandwidth, leading to minimal distortion in the resulting images. In this study, the bandwidth of the diffusion scans (393 Hz/Pixel) (18) is similar to the TSE scans (166 Hz/Pixel), requiring only a 6 degrees of freedom transformation to coregister the data.

To perform the fitting, T_2_ estimates from all white matter voxels were binned according to concentration (100 bins, range 0 – 1 for the KI and KT models, 0 – 23 mm for the D2S model), and the mean T_2_ estimated for each bin. Outliers (T_2_ estimates greater/less than the median ± 3 · median absolute deviation across all white matter) were not included in this calculation (and will not be included when presenting results in this manuscript). In very close proximity to the brain surface, T_2_ values were higher and characterised by a larger T_2_ error in comparison to other tissue. To avoid these boundary effects, voxels within 2 mm to the brain surface were additionally not included in these calculations. The binned data across the concentration range was fit to Eq. [5], with the fitting weighted by the number of voxels per bin.

The voxelwise influence of fixative concentration was subsequently eliminated over the entire brain to generate *T*_2_0%__(*x,y,z*) maps via:

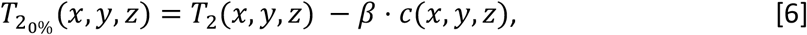

where *β* is a single brain-wide scalar. For the D2S model, *c*(*x, y, z*) is substituted for the distance to surface measurement.

In the absence of a ground truth, we require a metric for comparing across models. Although T_2_ is likely to vary across the brain within a given tissue type, spatial patterns matching a spatial model of fixative concentration should most conservatively be attributed to fixative. The fact that a single regression coefficient was fit to all of white matter means that it is unlikely to result in over-fitting. Therefore, performance of the different models was evaluated by comparing the homogeneity of the T_2_ maps before and after correction within tissue type. A concentration model is deemed to be “better” if it improves homogeneity (i.e. if it removes more variance) compared to another model.

### Correlation with Ferritin and PLP

This work forms part of a larger post-mortem imaging project investigating how changes in MR image contrast due to the neurodegenerative disease amyotrophic lateral sclerosis (ALS) relate to pathology as reflected in immunohistochemical staining (31). As part of this project, histological staining has been performed within each brain for ferritin (which is a non-quantitative surrogate for iron content in tissue) and PLP (a surrogate for myelin content in tissue). Tissue sections with these stains have been acquired in the primary motor cortex (M1), visual cortex (V2) and anterior cingulate cortex (ACC). Full details of the histology acquisition and processing pipeline are provided in (31). We assess the correlation between T_2_ and ferritin/PLP with and without correction for fixative concentration.

Histology results for PLP and ferritin are presented in terms of a stained area fraction (SAF). The SAF is defined as the ratio of the positively stained region of the analysed region of interest (ROI) relative to the total ROI. In this study, PLP SAF estimates are available for both hemispheres of M1 (in the leg, hand and face areas), V2 and the ACC. For ferritin, SAF estimates are available in the left brain hemisphere only for M1 (leg and face regions), V2 and the ACC. Ferritin staining was performed in two separate batches (Round 1 – M1 leg, V2 and ACC; Round 2 – M1 face, V2 and ACC). To account for cross-batch variability, each round was normalised (demeaned and divided by the standard deviation) prior to combination. To make comparisons with the T_2_ estimates, ROIs were generated in the diffusion space of the MRI data in the left and right hemispheres of M1, V2 and the ACC. For the motor cortex, standard space label masks were coregistered into the space of the post-mortem brains using FLIRT (22), followed by manual segmentation into leg, hand and face areas of the motor cortex. For V2 and ACC, masks were hand drawn in the space of the diffusion MRI data using the histology images as a guide. All masks were generated by a researcher familiar with neuroanatomy. Masks were subsequently coregistered into the space of the T_2_ maps using FLIRT (22). Any white matter areas were removed from the resulting masks prior to analysis and the T_2_ estimate was taken as the median value over the ROI.

## Results

Figure 5 displays the simulated outflux of fixative using the KI and KT models, alongside the phenomenological D2S model, in a single brain sample. Whereas the D2S model (Fig. 5c) reveals a markedly different distribution across the brain, only relatively subtle differences are observed between the KI (Fig. 5a) and KT (Fig. 5b) models. By taking the difference between these two maps (Fig. 6), it becomes apparent that the KT model exhibits an increased concentration of fixative in white matter and a decreased concentration of fixative in grey matter. This is consistent with a higher diffusion coefficient in grey matter vs the mean diffusivity, and a lower diffusion coefficient in white matter vs mean diffusivity, as can be inferred from (Fig. 1a).

**Figure 5:**
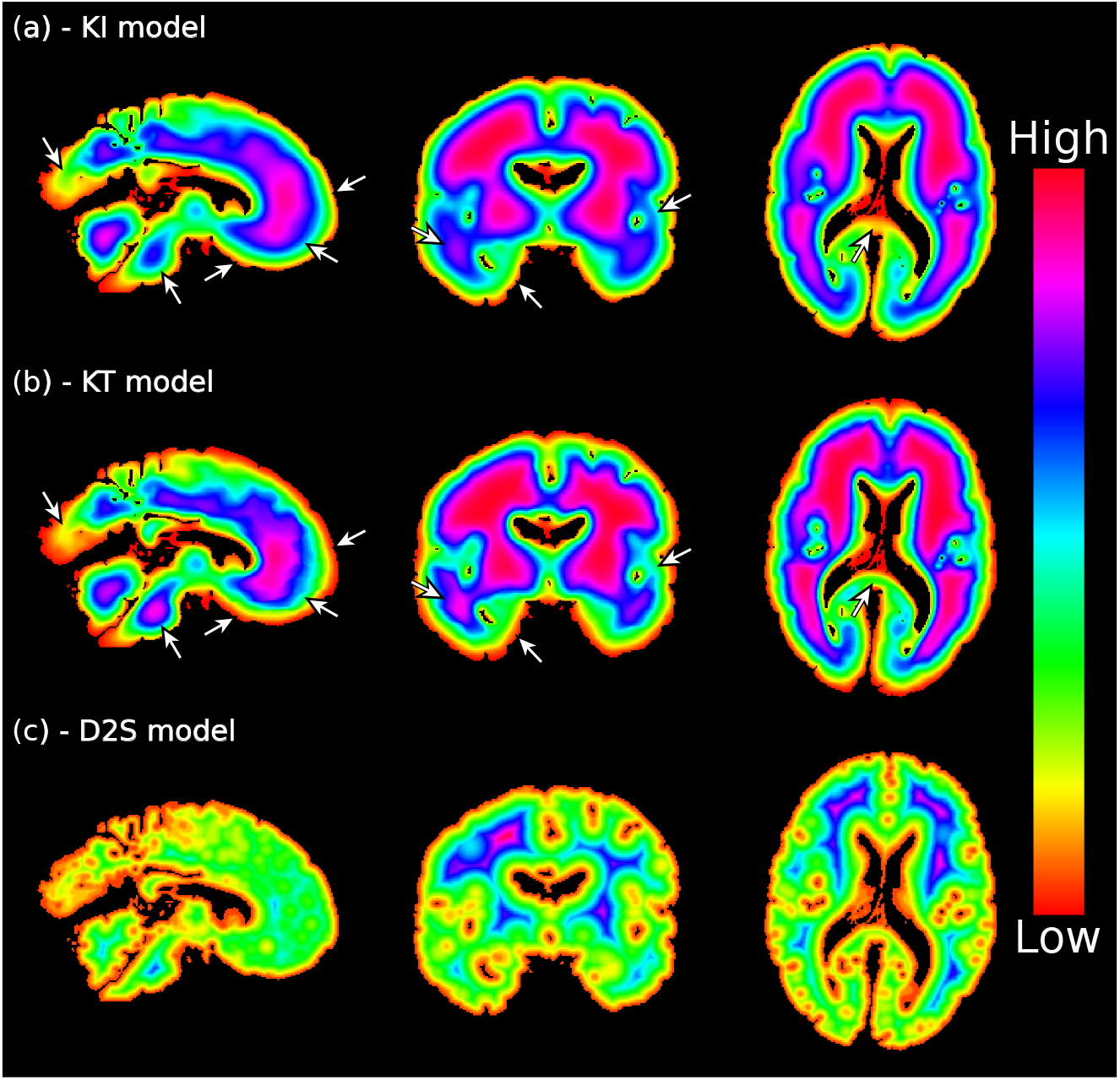
Modelling the outflux of fixative with the KI, KT and D2S model. Defining initial conditions from Eq. [4] (100% fixative concentration in tissue surrounded by an external medium of 0% fixative), here we display the resulting concentration distribution map for the kinetic isotropy (a) and kinetic tensor (b) models, and the phenomenological distance-to-surface (c) model. Subtle differences between the KI (a) and KT (b) models are apparent across both grey and white matter (white arrows). The D2S model (c) reveals a considerably different distribution across the brain. (a) and (b) modelled using the diffusion tensor estimates in Fig. 1 assuming fixative outflux for 48 hours. (a) and (b) are scaled between 0 and 1, with (c) scaled between 0 and 19.5 mm. Colormap chosen to highlight the differences across the brain.

**Figure 6:**
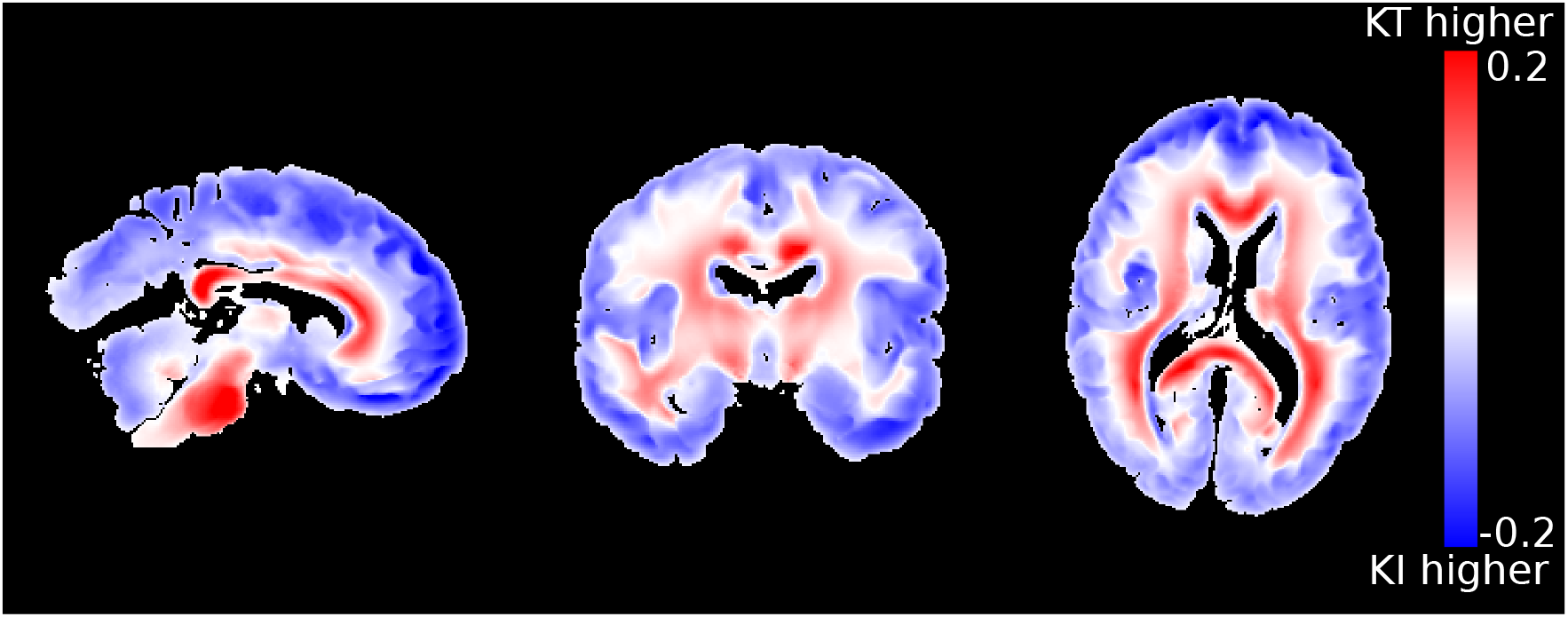
Differences between the KI and KT model. By examining the concentration difference between the KT and KI models (KT model minus KI model), it is apparent that the KT model is characterised by an increased fixative concentration across white matter, with a decreased concentration across grey matter vs KI. This is consistent with observations of an increased/decreased diffusion coefficient across grey/white matter in post-mortem brains (Fig. 1a) vs the mean diffusivity. Figure formed from the data in Fig. 5. Concentration distributions modelled using the diffusion tensor estimates in Fig. 1 assuming fixative outflux for 48 hours.

Figure 7 displays a single coronal slice of the T_2_ maps from all 14 brains, demonstrating that our EPG framework (details provided in Supporting information) generates T_2_ maps that exhibit consistent contrast across grey and white matter for each fixative type. Figure 7 additionally reveals that the fixative type has a considerable effect on the magnitude of T_2_ estimates, with an increased T_2_ observed in both white and grey matter (Fig. 8) for brains fixed with 10% NBF vs 10% formalin. No significant associations were found between the mean T_2_ across the entire brain and the post-mortem delay / time in fixative before scanning (values provided in Supporting Information Table S1) for each fixative type.

**Figure 7:**
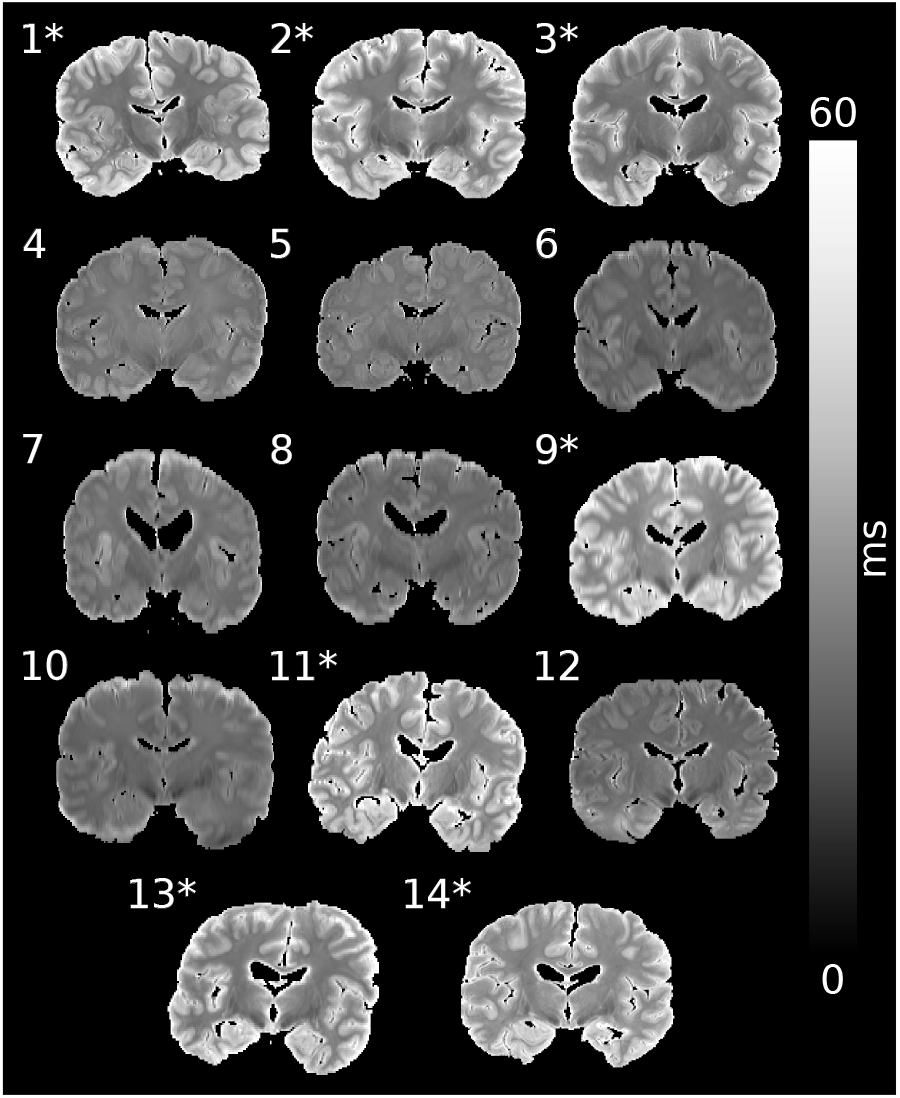
Single coronal slice of the T_2_ maps from all 14 brains. Our EPG framework (details provided in Supporting Information) accounts for the influence of B_1_ homogeneity at 7T, reducing the bias on T_2_ estimates in areas of low B_1_. Brains fixed with 10% NBF (*) display significantly higher T_2_ estimates in both white and grey matter (see Fig. 8).

**Figure 8:**
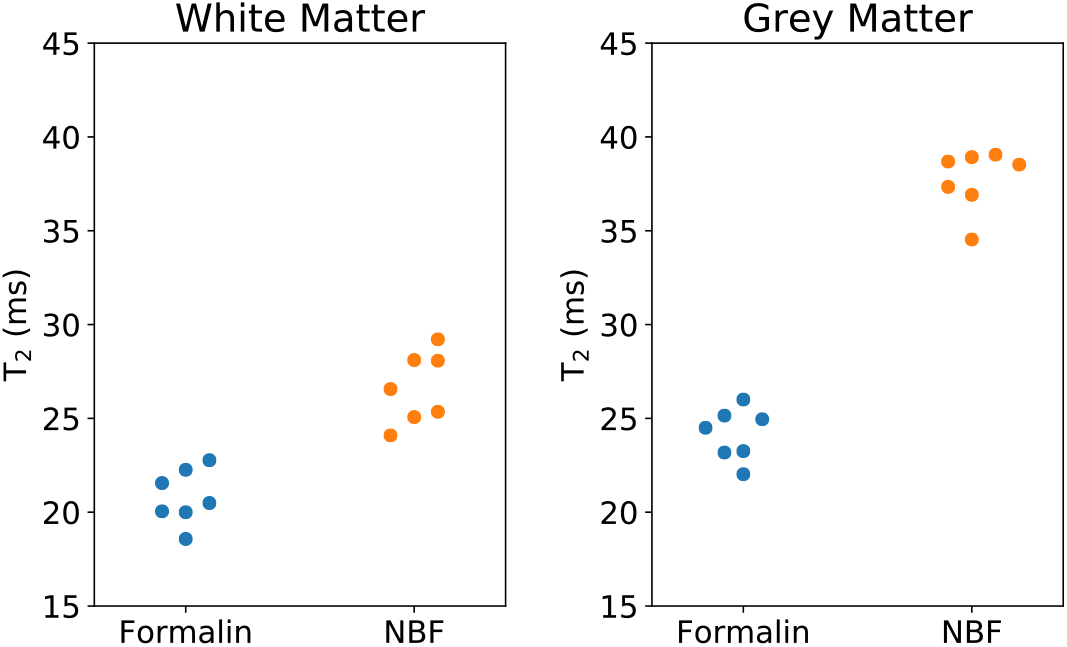
Mean T_2_ for brains fixed with 10% formalin and 10% NBF over white and grey matter. Brains fixed with 10% NBF were characterised by a higher estimate of T_2_ over white matter (p = 4.2 · 10^−5^, Cohen’s D = 3.2) and grey matter (p = 1.4 · 10^−9^, Cohen’s D = 8.3), with differences clearly depicted in Fig 7. Each dot represents the mean T_2_ over white/grey matter for a single brain. p-values estimated using Welch’s t-test. Horizontal displacement along x-axis for individual points for visualisation purposes only.

Figure 9 displays the relationship of T_2_ vs concentration across white and grey matter using the D2s, KI, and the KT models for brains fixed with NBF. In all cases, the model appears to explain a large amount of variation in T_2_. A linear decrease in T_2_ with increases in fixative concentration is apparent for the KI and KT models prior to correction (Fig 9a), in agreement with previous reports (4). The D2S model similarly displays a decrease in T_2_ with increased distance to surface, but is more inhomogeneous across the distance profile. In addition, the binning of voxels according to the D2S model results in higher standard deviation, suggesting that distance to surface is less relevant to predicting a voxel’s T_2_ than the KI and KT concentration models. By correcting for the influence of fixative concentration using Eq. [6] (Fig. 9b), all three models produce flatter profiles across a wide range of concentrations in white matter (i.e. voxels included in the fit), and reduce the inhomogeneity across grey matter (i.e. voxels not include in the fit). The KI and KT models produced notably flatter profiles compared to D2S. Interestingly, brains fixed with formalin did not show the same trend, with changes in T_2_ on the order of a few ms over the entire concentration range (Fig. 10). Correction across these samples led to little observable change for all three models.

**Figure 9:**
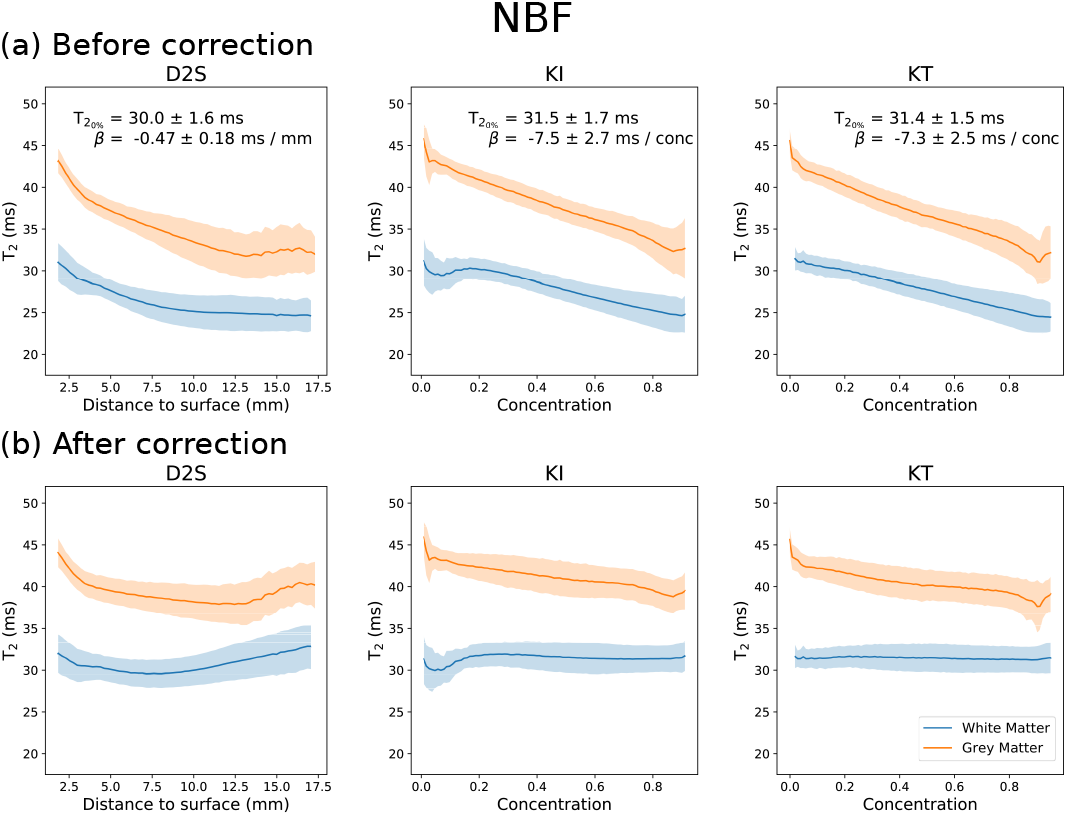
T_2_ vs concentration/distance to surface over white and grey matter for all post-mortem brains fixed with 10% NBF. Averaging over all brains fixed with 10% NBF, all three models display a decrease in T_2_ with increased concentration/distance to surface (a). Whereas the KI and KT models demonstrate a linear relationship (in agreement with (4)), the D2S model displays a more inhomogeneous change in T_2_. Regressing out the influence of fixative concentration using Eq. [6] improves the homogeneity of T_2_ estimates across white and grey matter in all three models (b). Results displayed as the mean ± standard deviation across all brains fixed with 10% NBF.

**Figure 10.**
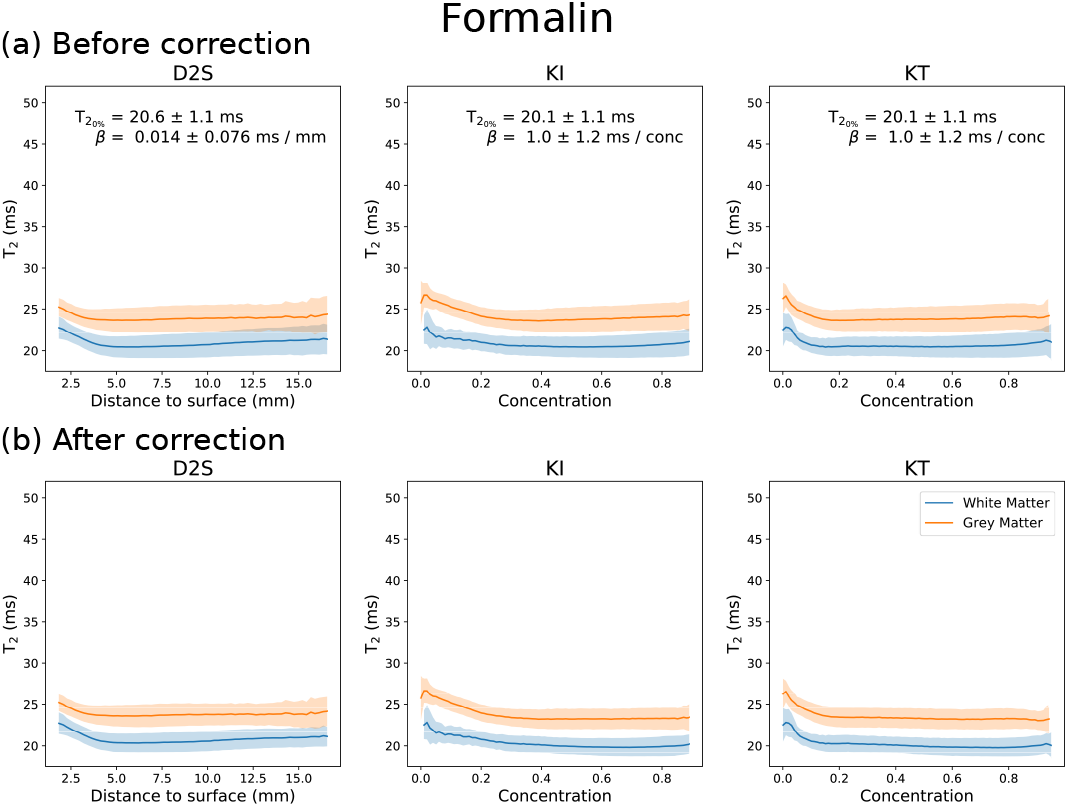
T_2_ vs concentration/distance to surface over white and grey matter for all post-mortem brains fixed with 10% formalin. Averaging over all brains fixed with 10% formalin, all three models display a small decrease in T_2_ with increased concentration/distance to surface (a). This change is inhomogeneous across all three models, where the change in T_2_ is characterised by a small *β* for all three models. Regressing out the influence of fixative concentration using Eq. [6] leads to little apparent change across white and grey matter in all three models (b). Results displayed as the mean ± standard deviation across all brains fixed with 10% formalin.

Table 1 displays the inhomogeneity (defined here in terms of the standard deviation) across brains fixed with 10% NBF (Table 1a) and 10% formalin (Table 1b) within grey and white matter separately before and after correction. T_2_ maps for brains fixed with 10% NBF are characterised by a higher inhomogeneity across both white and grey matter prior to correction. In these brains (Table 1a), corrections based on all three models reduced inhomogeneity. Notably, this improvement is observed for both white and grey matter, despite the model being fit to white matter voxels only. The KI and KT models reveal similar performance, with the KT model demonstrating the best overall improvement (lowest inhomogeneity over both white and grey matter). Over white matter, the reduction in inhomogeneity reaches significance (defined as p < 0.05) for the KI and KT models. Over grey matter, although none of the models reach significance, the KT model is very close and demonstrates the best overall reduction in inhomogeneity. Figure 11 displays a 10% NBF brain before and after correction, demonstrating a visible reduction in inhomogeneity across the brain.

**Figure 11:**
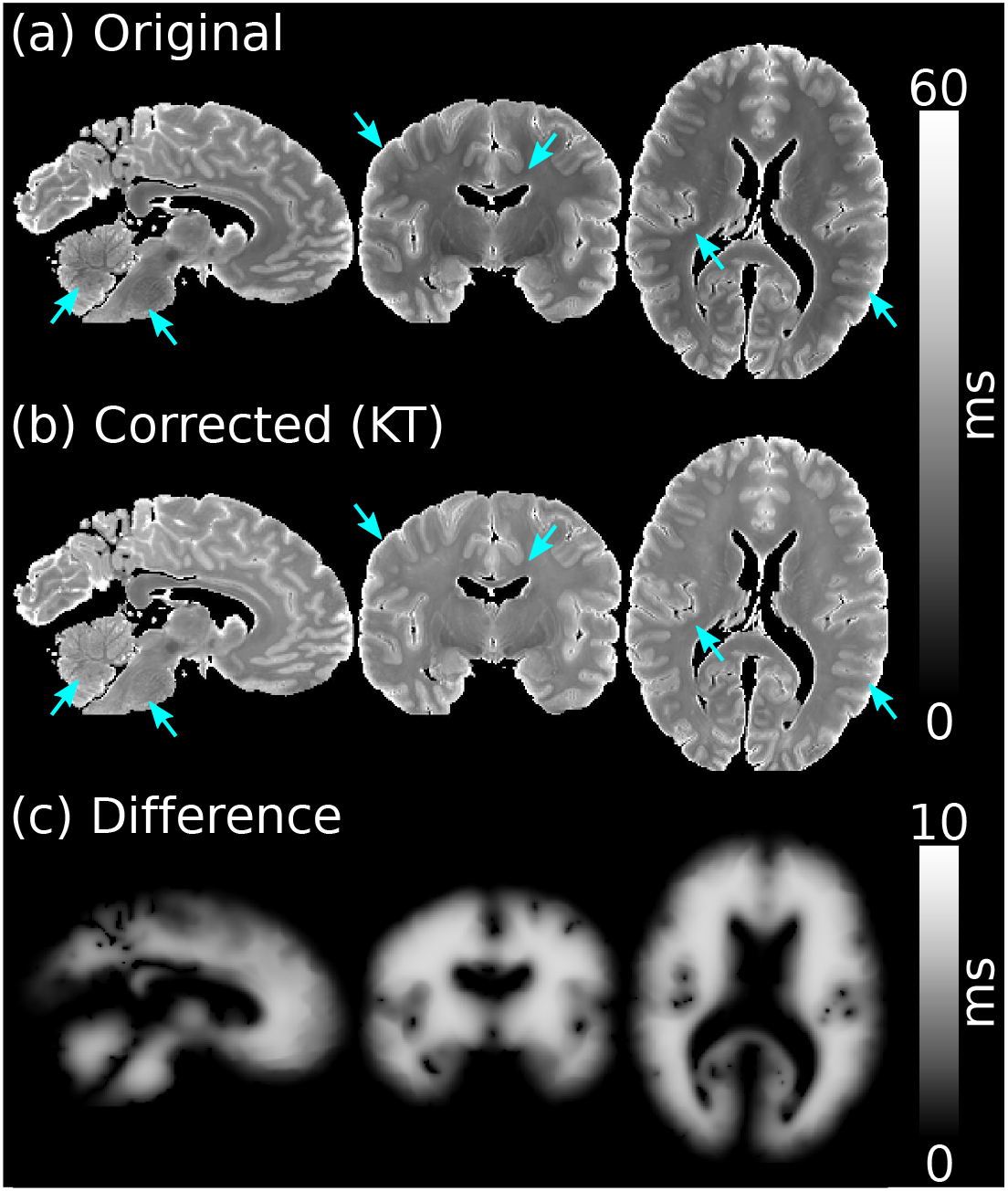
T_2_ map before and after correction with the KT model. By performing a correction with the KT model over a postmortem T_2_ map (a), we are able to reduce the inhomogeneity across the brain (b). These differences are most apparent within regions close to the brain surface (a and b arrows). The difference map (c – corrected minus original) is a scaled KT concentration distribution.

**Table 1:**
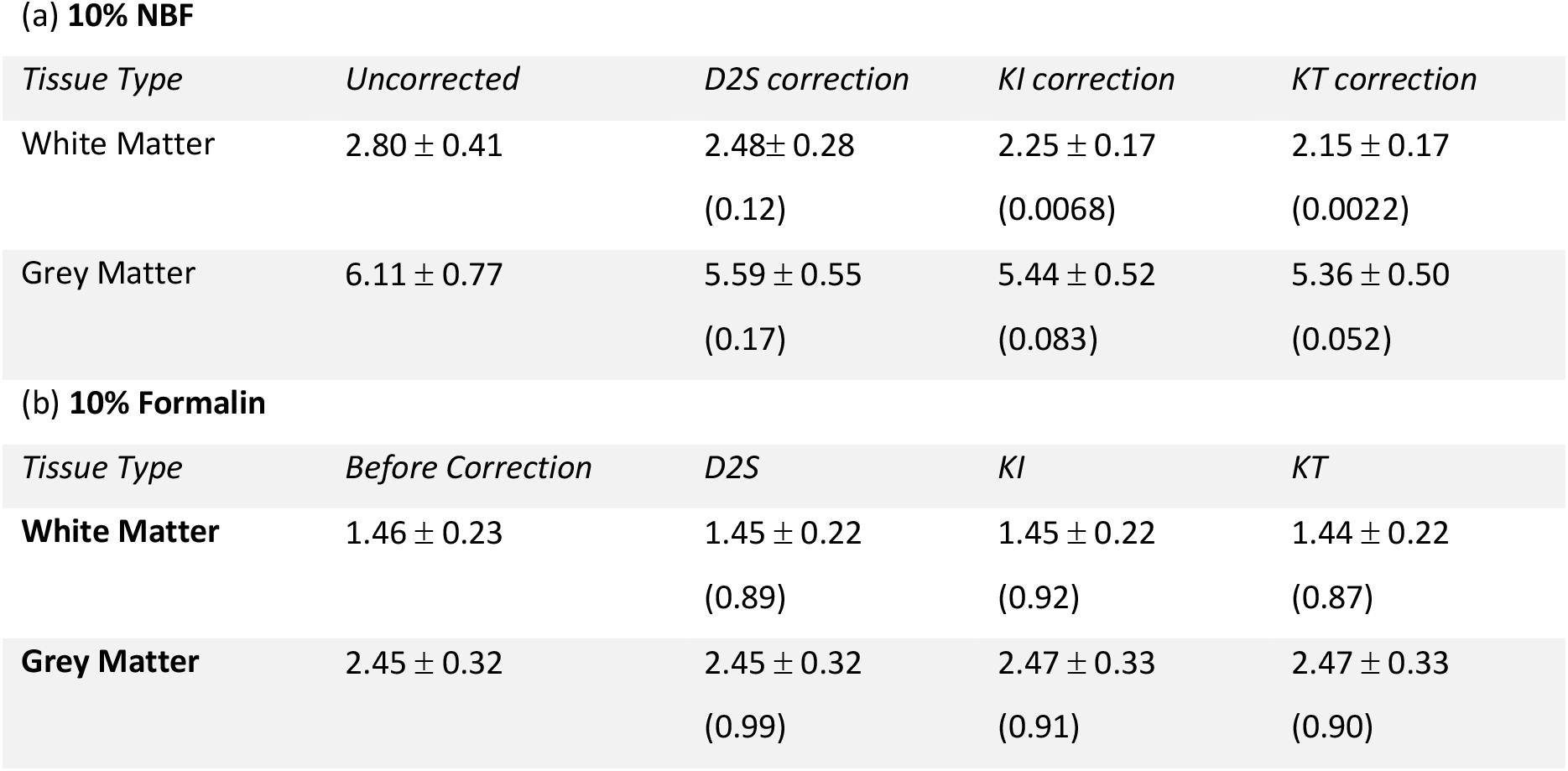
Inhomogeneity across white and grey matter for brains fixed with 10% NBF and 10% formalin. For brains fixed with 10% NBF (a), all three models lead to a reduction in inhomogeneity (defined here as the standard deviation) across the brain. The KI and KT models generate a reduced inhomogeneity across both white and grey matter vs the D2S model. The KI and KT models perform similarly, with the KT model demonstrating the best overall improvement. For the brains fixed with 10% formalin (b), all three models lead to very little change in inhomogeneity, notably an insignificant increase in inhomogeneity with the KI/KT models across grey matter.

**(a) 10% NBF**
| Tissue Type | Uncorrected | D2S correction | KI correction | KT correction |
| --- | --- | --- | --- | --- |
| White Matter | $2.80 \pm 0.41$ | $2.48 \pm 0.28$<br>(0.12) | $2.25 \pm 0.17$<br>(0.0068) | $2.15 \pm 0.17$<br>(0.0022) |
| Grey Matter | $6.11 \pm 0.77$ | $5.59 \pm 0.55$<br>(0.17) | $5.44 \pm 0.52$<br>(0.083) | $5.36 \pm 0.50$<br>(0.052) |

**Table 1: Inhomogeneity across white and grey matter for brains fixed with 10% NBF and 10% formalin.**
| Tissue Type | Before Correction | D2S | KI | KT |
| --- | --- | --- | --- | --- |
| White Matter | $1.46 \pm 0.23$ | $1.45 \pm 0.22$<br>(0.89) | $1.45 \pm 0.22$<br>(0.92) | $1.44 \pm 0.22$<br>(0.87) |
| Grey Matter | $2.45 \pm 0.32$ | $2.45 \pm 0.32$<br>(0.99) | $2.47 \pm 0.33$<br>(0.91) | $2.47 \pm 0.33$<br>(0.90) |

For brains fixed with 10% formalin, none of the corrections lead to much difference in inhomogeneity across both white and grey matter (Table 1b), characterised by very small changes which do not reach significance. In these brains, the KI and KT models lead to a small increase in inhomogeneity across grey matter (possible when considering the regression parameters are estimated using white matter only).

Comparisons with histology reveal a negative correlation between T_2_ and PLP for both the 10% NBF and formalin brains (Fig. 12). Brains fixed with 10% NBF demonstrate a stronger negative correlation than those fixed with 10% formalin, with the correlation for brains fixed with 10% formalin just below significance. Correction with the KT model increased the similarity between the two fixative types, with a small decrease in the correlation between T_2_ and PLP for brains fixed with 10% NBF, and a small increase for brains fixed with 10% formalin (reaching significance after correction). For the ferritin results (Fig. 13), a small negative correlation was found for brains fixed with 10% NBF, which was reduced (and lost significance) after correction with the KT model. No significant correlation found for brains fixed with 10% formalin before or after correction.

**Figure 12:**
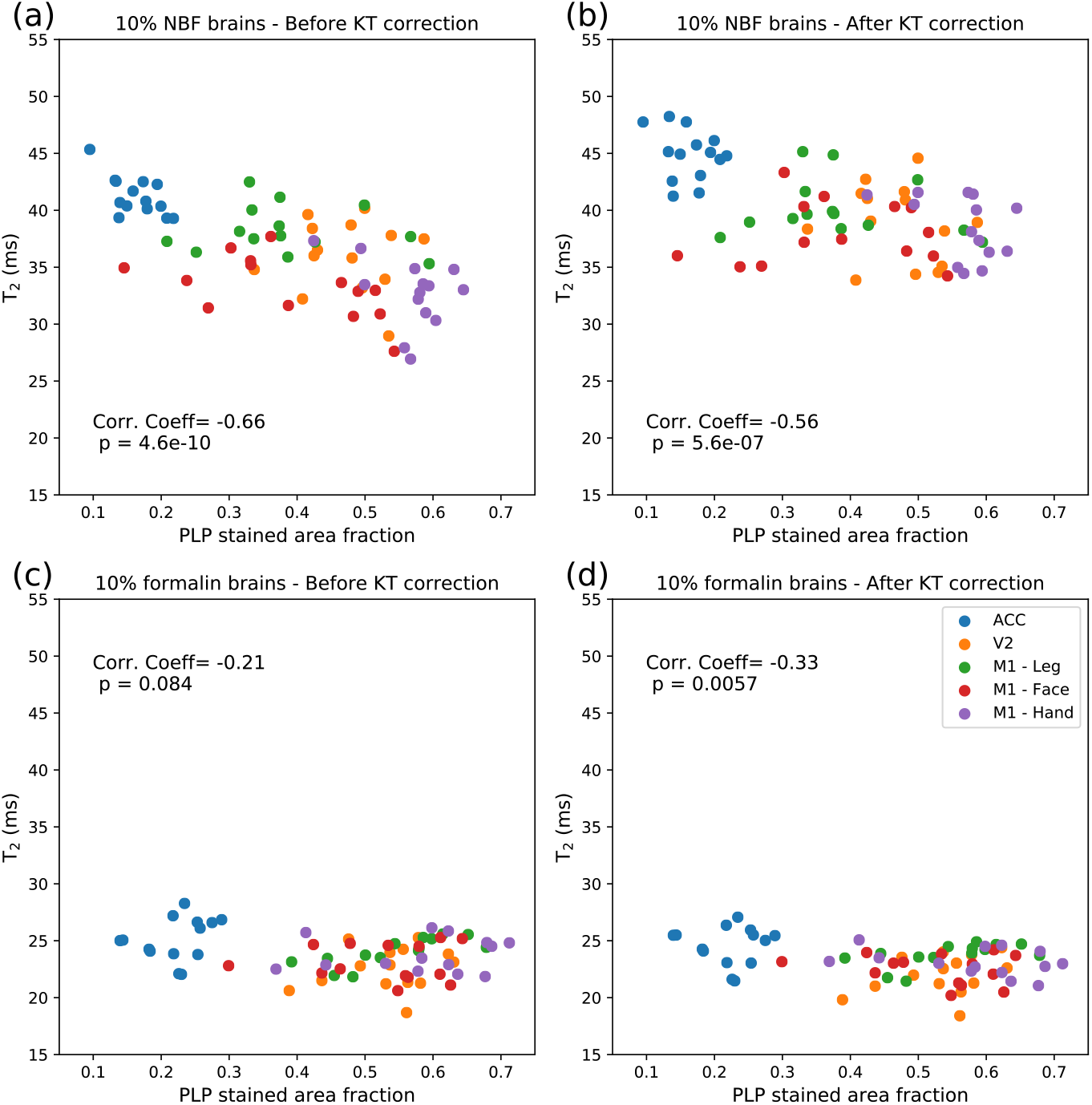
Correlation between T_2_ and PLP for brains fixed with 10% NBF and 10% formalin. Brains fixed with 10% NBF (a) and 10% formalin (c) demonstrate a negative correlation with PLP, with the relationship predominantly driven by regional differences in PLP/T_2_ across the ROIs used in this study. Correction with the KT model (b and d) improved the similarity of the relationship between the two fixative types, corresponding to a reduced/increased correlation for brains fixed with 10% NBF/10% formalin respectively.

**Figure 13:**
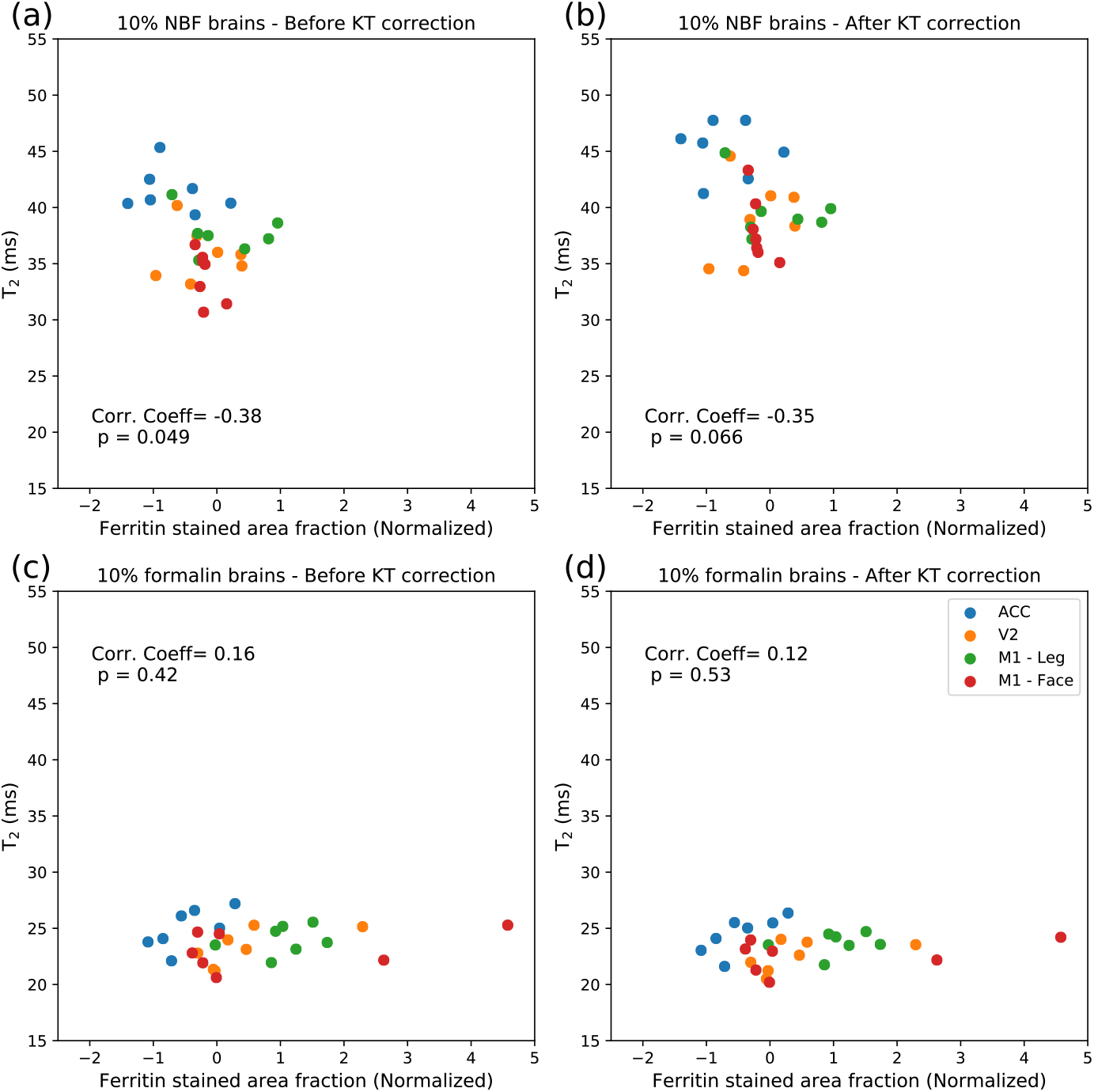
Correlation between T_2_ and ferritin for brains fixed with 10% NBF and 10% formalin. Brains fixed with 10% NBF (a) display a small negative correlation with ferritin, where correction, with a reduced correlation (and a loss of significance) after KT correction. For brains fixed with 10% formalin, no correlation was observed before (c) or after (d) the KT correction. Note that as the ferritin SAFs were normalised for the two batches, the SAF values can be positive & negative and are not restricted to a range between 0 and 1. As the ACC and V2 regions were included in both batches, the ferritin SAFs were averaged prior to plotting.

## Discussion

In this study, we have expanded on existing literature for modelling fixative dynamics. The KI model, which assumes a single brain-wide diffusion coefficient and models the effect of geometry on the influx of fixative, is based on the work by Dawe et al. (10). We introduced the KT model, which incorporates the effects of diffusion anisotropy and tissue specific diffusion coefficients. Our results reveal that brains fixed with 10% NBF were found to have a strong dependence on profiles of fixative outflux, whereas brains fixed with 10% formalin were not found to have such a dependence. Models that incorporate realistic fixative dynamics (KI and KT) were found to yield T_2_ maps with a reduced inhomogeneity vs a simple distance to surface model, with the greatest reduction in inhomogeneity attributed to the KT model.

Brains fixed with 10% NBF and 10% formalin were found to be characterised by very different T_2_ properties, with brains fixed with 10% NBF generating higher overall T_2_ estimates in both grey and white matter (Fig. 8). This observation highlights the importance of accounting for the specific composition of the formalin solution when analysing post-mortem data, where the choice of buffer has a considerable influence on T_2_ (even when considering the same formalin concentration). Previous work (6) has observed that even the vendor-specific composition of the fixative solution is a substantial contributor to the estimated MR relaxation properties. What is more surprising here is the distinction between the two fixative types in T_2_ with respect to fixative outflux, given that the only difference between the two fixatives should be the buffer solution. This result suggests that if we are measuring a change in fixative concentration due to outflux at the brain surface, the composition of the fixative solution may lead to a more complicated relationship with the estimated T_2_. However, as no external validation was performed of the fixative outflux over the course of this experiment, this hypothesis cannot be tested further.

Comparisons with histology reveal that brains fixed with 10% NBF demonstrate an overall stronger correlation with both PLP and ferritin vs brains fixed with 10% formalin (Figs. 12 and 13). For the PLP analysis, brains fixed with 10% NBF and 10% formalin both demonstrate a negative correlation (Fig. 12), consistent with the observation that myelin is characterised by a short T_2_, and an increased myelin content leads to a decrease in T_2_ (35,36). The correlation appears to be predominantly driven by regional differences, with the ACC characterised by the lowest level of myelination vs V2 and M1 (in agreement with (37,38)), and T_2_ estimates broadly reflecting these SAFs.

Correction with the KT model increased the similarity between the relationship of T_2_ with PLP for the two fixative types. By correcting for the concentration of fixative with the KT model, we reduce the variance of T_2_ across different regions of the brain within individual subjects. This correction lead to a small decrease in the correlation with PLP for brains fixed with 10% NBF, suggesting that the confound of fixative concentration is artificially inflating the correlation between T_2_ and PLP in these brains. Although we would typically expect the removal of confounds to increase the correlation between regions under these conditions, as the correlation between T_2_ and PLP is predominantly driven by regional differences, any regional dependencies on fixative concentration could drive this correlation. A small (but significant) positive correlation was found between the expected fixative concentration (as simulated by the KT model) and PLP SAF (Supporting Information Fig. S5). This suggests that if is there is outflux of fixative at the tissue surface (which leads to a characteristic change in T_2_), then the correlations across different brain regions are partially driven by the fixative concentration. No significant correlation between concentration and PLP SAF was found for brains fixed with 10% formalin (Supporting Information Fig. S6), where a small increase in correlation was observed after correction.

The correlations between T_2_ and ferritin SAF are low, with only a small negative correlation reached for brains fixed with 10% NBF (losing significance after correction) and no notable correlation for brains fixed with 10% formalin. In addition, no significant correlation was found between fixative concentration and ferritin SAF for either fixative type (Supporting Information Fig. S6). Although ferritin is a non-quantitative estimate of tissue iron, we would expect an increased ferritin content to correspond to a decrease in tissue T_2_ (39). However, there are a number of limitations to our ferritin analysis that could explain this low level of correlation, most notably that ferritin staining is highly variable between batches and staining quality is very operator dependent. Although some effort was taken to normalise the results and combine across batches, when combined with the limited number of regions where ferritin staining data is available makes us particularly sensitive to outliers. Further details of these limitations have been described in detail in a recent publication from our group (21). We are currently exploring alternative approaches to more accurately quantify the ferritin content of tissue and move away from simple summary statistics provided by SAFs, most notably with the development of a toolbox to directly coregister histology slides to MRI images (40). This will enable us to perform more sophisticated voxelwise comparisons between the MRI and histology data.

There are several limitations to this study. First, no external validation of the outflux of fixative from the post-mortem brain sample was performed. Therefore, whilst we observe a correlation between our concentration distribution and the T_2_ estimates in brains fixed with 10% NBF, we cannot confirm that this is due to fixative outflux. Although correction with the KT model does appear to remove inhomogeneity in these brains (e.g. Fig. 11 and Table 1), the inconsistencies between the two fixative types remains unexplained and requires further exploration. Second, the KI/KT simulations have a strong dependency on the outflux duration. The duration of time between the brains being placed in fluorinert and scanning was not accurately recorded for each individual sample, with 48 hours chosen as an approximate time between these two events. However, our simulations additionally reveal that the concentration distribution does not evolve linearly with time (Fig. 14). Precise knowledge of this time period is recommended for accurately simulating the effects of fixative outflux. Similarly, the choice of b-value in the diffusion MRI experiment may lead to different diffusivity estimates in the post-mortem tissue sample (due to non-Gaussian diffusion within tissue (41)) and thus differences in the concentration profile. Third, the TSE sequence used in this study was highly sensitive to B_1_, requiring the use of an EPG fitting approach to estimate our T_2_ maps (detailed in Supporting Information). We additionally investigated whether any of the observed correlations could be attributed to the B_1_ distribution, which has a similar spatial profile to the outflux models used in this study. Although regressing out the B_1_ distribution did lead to a decrease in inhomogeneity (Supporting Information Fig. S7 and Table S3), this decrease in inhomogeneity was lower in comparison to the D2S, KI and KT model over brains fixed with 10% NBF, and similar in performance for brains fixed with 10% formalin (where the concentration correction did not lead to any significant change in inhomogeneity).

**Figure 14:**
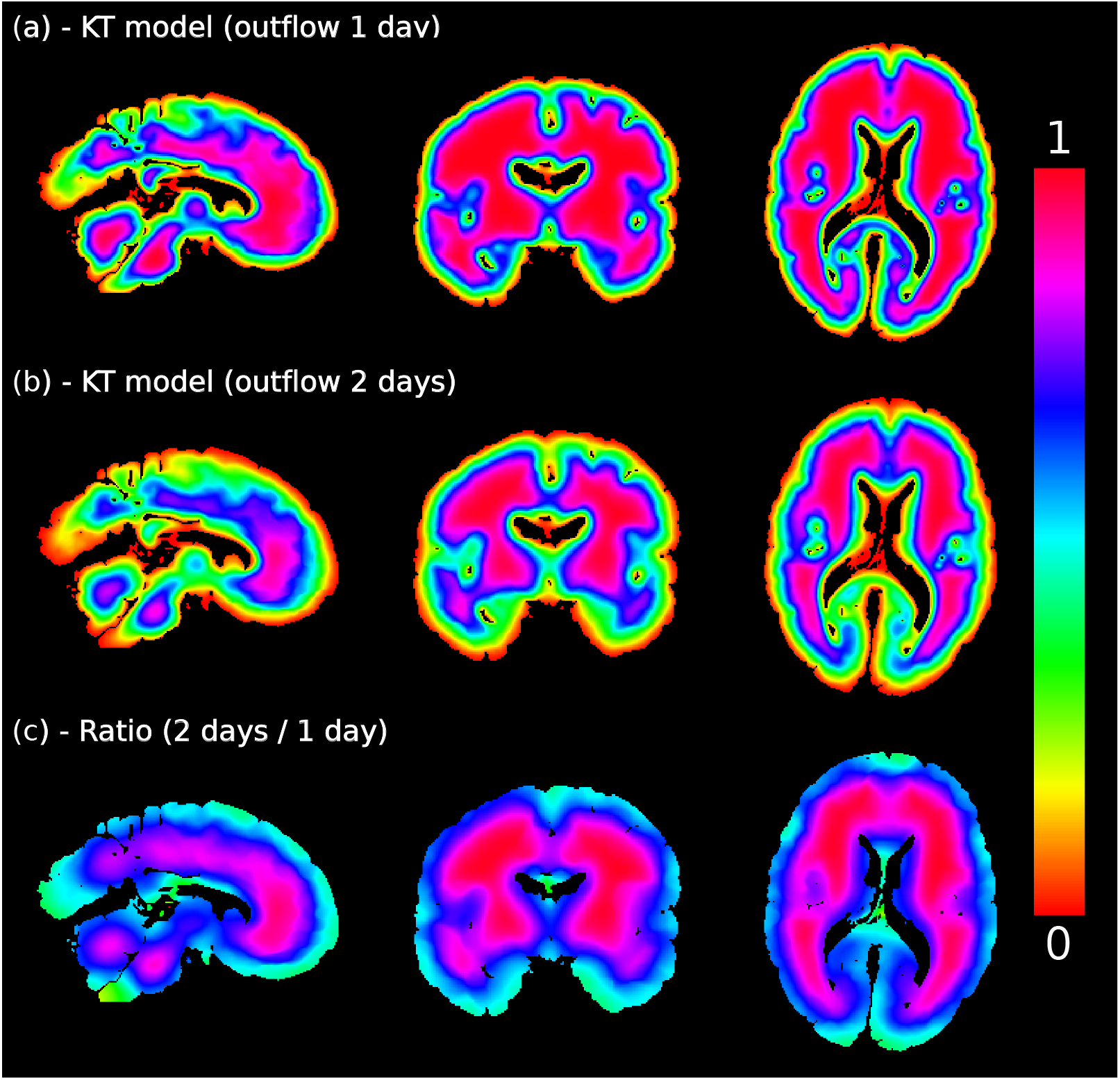
Non-linearity of the KT model. Here we display simulations of outflux for one (a) and two (b) days using the KT model. The concentration distribution across the brain does not scale linearly with time. This leads to a ratio map (c) that does not reflect the same value across the entire brain. It is therefore recommended to have precise recordings of influx/outflux duration in order to use this approach effectively.

Application of the KT model in this study used diffusion tensor estimates acquired in the same post-mortem brain to model fixative dynamics. However, for post-mortem studies that do not perform diffusion MRI as part of their acquisition, the use of a diffusion-tensor atlas (e.g. the HCP0165 standard space DTI template, available in FSL (33)) could be explored as an alternative approach. The KI model (which additionally demonstrated improved performance vs the phenomenological D2S model) requires only a mask of the tissue sample and a single estimated mean diffusivity to simulate.

This work forms part of a larger project (31) investigating the pathology of ALS through the combination of post-mortem MRI and immunohistochemical staining within the same tissue sample, to determine how changes in tissue composition gives rise to measured changes in our MR signal. In order to accurately map these relationships, it is essential to remove any potential confounds which could mask out subtle changes in the MR signal due to tissue pathology, or drive spurious relationships in our data. In this manuscript, we focused on using the KT model to correct for fixative concentration due to the outflux of fixative. However, it would be possible to extend this approach to other challenges in postmortem imaging. One example is the estimation of a voxelwise post-mortem delay. When a brain is fixed, fixative penetrates slowly into brain tissue (Fig. 2). By modelling the influx of fixative into tissue, it would be possible to generate a voxelwise estimate of the time required for any individual voxel to become fully fixed. This could additionally be modelled and removed as a confound in the data. A voxel-wise post-mortem delay (42) might be predictive of effects related to cross-linking of tissue, which in turn may be reflected in MRrelevant properties like T1.

## Conclusion

We have introduced the KT model, which incorporates diffusion anisotropy and tissuespecific diffusion properties when modelling fixative dynamics within tissue. By modelling fixative dynamics in tissue, we have additionally demonstrated that the resulting map can be used to remove confounds from MR images. T_2_ maps acquired in whole post-mortem brains reveal a spatial profile consistent with a model of fixative outflux in brains fixed with 10% NBF, with the KT model yielding the greatest reduction in inhomogeneity in T_2_ across both grey and white matter. Results were found to be strongly dependent on the type of fixative, with further exploration required to determine whether the observed changes can be attributed to fixative outflux, and the contribution of the buffer solution to this process.

## Acknowledgements

This study was funded by a Wellcome Trust Senior Research Fellowship (202788/Z/16/Z) and a Medical Research Council grant (MR/K02213X/1). Brain samples were provided by the Oxford Brain Bank (BBN004.29852). The Wellcome Centre for Integrative Neuroimaging is supported by core funding from the Wellcome Trust (203139/Z/16/Z). We acknowledge the Oxford Brain Bank, supported by the Medical Research Council (MRC), Brains for Dementia Research (BDR) (Alzheimer Society and Alzheimer Research UK), and the NIHR Oxford Biomedical Research Centre. The views expressed are those of the authors and not necessarily those of the NHS, the NIHR or the Department of Health.

## Appendix

Eq. [1] is discretised over space and time to obtain (19):

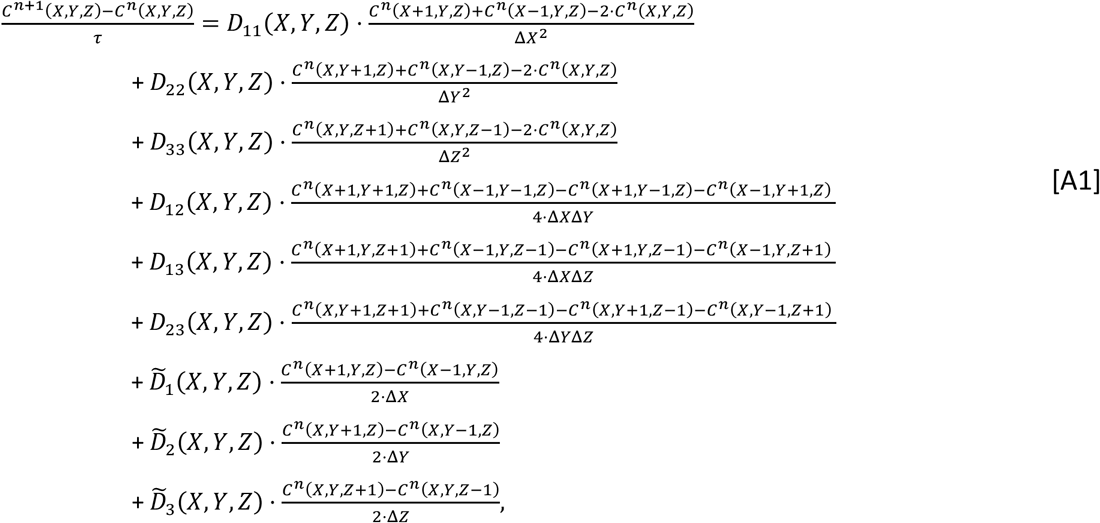

where:

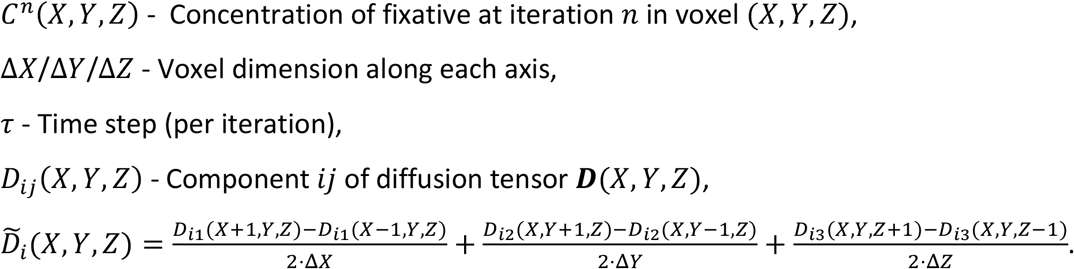

By rearranging Eq. [A1], the spatial distribution of fixative concentration at iteration *n* + 1 (*C*^*n*+1^) can be estimated from *C^n^* and ***D***.

## Notes

### Competing Interest Statement

The authors have declared no competing interest.

