## Supporting Information for "A method to remove the influence of fixative concentration on post-mortem T_2_ maps using a Kinetic Tensor model"

| Brain | Clinical diagnosis | Post-mortem delay (days) | Fixative type | Time in fixative before scanning (days) |
|---|---|---|---|---|
| **1** | Control | 3 | 10% NBF | 115 |
| **2** | Control | 3 | 10% NBF | 45 |
| **3** | Control | 3 | 10% NBF | 48 |
| **4** | ALS | 4 | 10% Formalin | 178 |
| **5** | ALS | 3 | 10% Formalin | 114 |
| **6** | ALS | 7 | 10% Formalin | 137 |
| **7** | ALS | 4 | 10% Formalin | 94 |
| **8** | ALS | 3 | 10% Formalin | 158 |
| **9** | ALS | 1 | 10% NBF | 35 |
| **10** | ALS | Data Unavailable | 10% Formalin | 96 |
| **11** | ALS | 2 | 10% NBF | 87 |
| **12** | ALS + FTD | 2 | 10% Formalin | 283 |
| **13** | ALS | 3 | 10% NBF | 94 |
| **14** | ALS | 2 | 10% NBF | 139 |

Table S1: **Characteristics of each brain used in this study**. Here ALS – Amyotrophic Lateral Sclerosis, FTD – Frontotemporal Dementia, NBF – Neutral Buffered Formalin.

| Brain | 1-3, 14 | 4 | 5-8 | 9-10 | 11-13 |
|---|---|---|---|---|---|
| **No. echoes** | 6 | 6 | 6 | 6 | 6 |
| **TE (ms)** | 13, 25, 38, 50, 63, 76 | 11, 23, 34, 46, 57, 69 | 11, 23, 34, 46, 57, 69 | 11, 23, 34, 46, 57, 69 | 10, 20, 30, 40, 50, 60 |
| **TR (ms)** | 1000 | 1000 | 1000 | 1000 | 1000 |
| **Resolution ($mm^3$)** | 0.9 x 0.9 x 0.9 | 0.8 x 0.8 x 1.6 | 0.8 x 0.8 x 1.6 | 0.65 x 0.65 x 1.3 | 1.0 x 1.0 x 1.2 |
| **Bandwidth (Hz/pixel)** | 166 | 195 | 195 | 163 | 199 |
| **Turbo factor** | 6 | 6 | 6 | 6 | 6 |
| **Time per TE (minutes)** | 36 | 26 | 16 | 18 | 24 |
| **Slice resolution (%)** | 100 | 100 | 50 | 50 | 100 |

Table S2: **Acquisition parameters for the TSE scans used in this study**. Details of individual brains provided in Table S1.

**Estimating $T_2$ using an Extended Phase Graph (EPG) model**

The TSE sequence is characterised by a train of RF pulses ($90^o$-$180^o$-$180^o$-$180^o$-…), where the signal is sampled after each $180^o$ pulse. Assuming a single tissue compartment and a perfect $180^o$ pulse, the TSE signal evolves via a characteristic mono-exponential decay (Fig. S1a – red line). At 7T, $B_1$ inhomogeneity produces a spatially varying flip angle across the brain (Fig. S1b), leading to refocusing pulses that vary from $180^o$. Under these conditions, the signal evolution can deviate substantially from a mono-exponential signal model (Fig. S1a – purple, blue and green lines). Fitting a mono-exponential signal model to these data will lead to estimates of $T_2$ that strongly depend on the $B_1$ profile (Fig. S1c).

Extended Phase Graphs (EPG) provide a more accurate description of signal evolution under a variety of MRI sequences and conditions (1–3). In this work, we use an EPG framework to describe the signal evolution of the TSE sequence under a refocusing pulse of flip angle $\alpha$. Our approach fits the measured TSE signal to estimate the voxelwise $T_2$ and $\alpha$ (via estimation of $B_1$). Fitting with an EPG model is shown to reduce the bias on $T_2$ estimates in areas of low $B_1$, leading to more homogeneous $T_2$ maps across the brain (Fig. S1d).

To achieve this, we performed a two-step fitting approach, where we first fit the TSE signal to an EPG model to estimate $T_2$ and $B_1$ across the brain. It was found that in regions of very low $B_1$, sharp discontinuities were observed in the $B_1$ maps. Given that $B_1$ is expected to smoothly vary across the brain, we subsequently smoothed the $B_1$ map and repeated the fitting for $T_2$, keeping $B_1$ as a fixed parameter.

*Code availability*

Code for the EPG fitting used in this study is available at https://github.com/BenjaminTendler/KT_model. This fitting is based on EPG software associated with (23), written by Matthias Weigel in MATLAB. This software is available via contacting Matthias at.

*Step one:*

For each postmortem brain we simulated the TSE sequence (parameters provided in Table S2) under an EPG framework, obtaining voxelwise estimates of $B_1$ and $T_2$ by minimising:

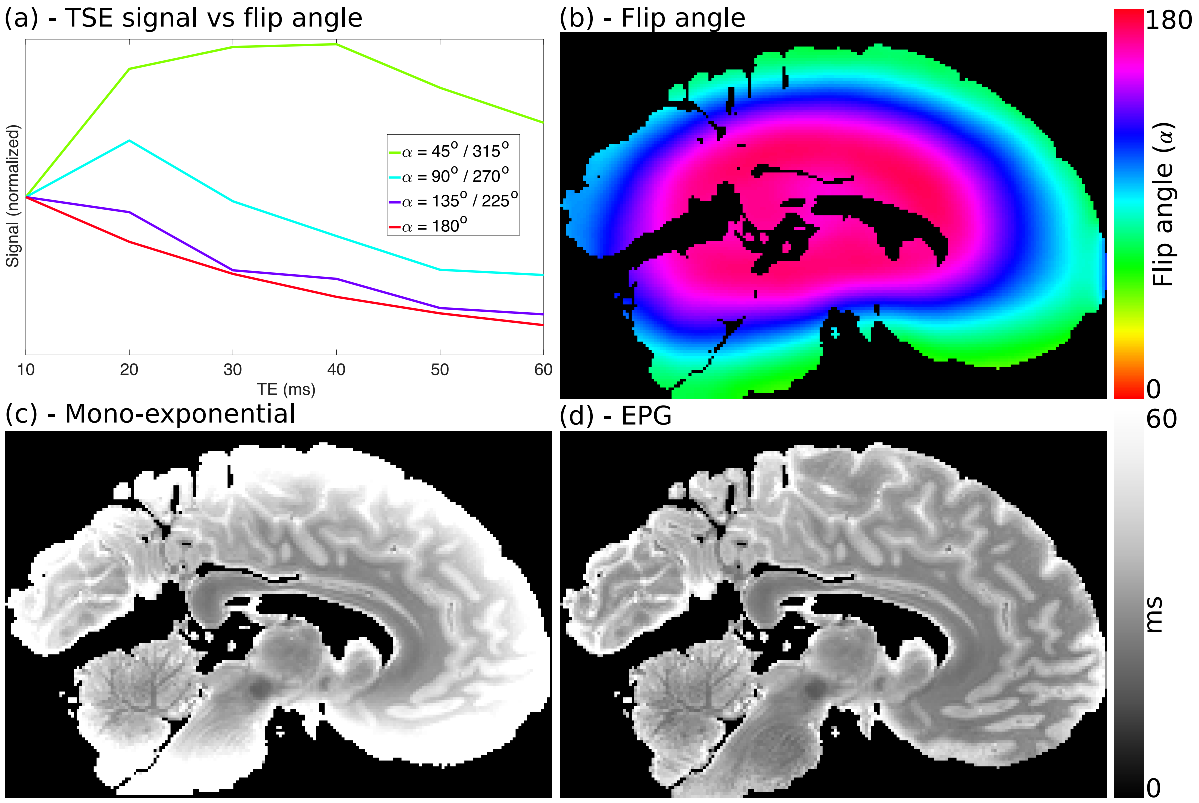


Figure S1: **Motivation for the EPG framework**. When the refocusing angle = 180° (a – red line), the TSE signal evolves via a mono-exponential decay. However, at different flip angles the signal evolution can deviate substantially from a mono-exponential signal model (a – purple, blue and green lines). At 7T, brain samples experience $B_1$ inhomogeneity, leading to a spatially varying flip angle across the brain (b). Fitting these data with a mono-exponential model leads to a $T_2$ map that depends strongly on the $B_1$ profile (c). An EPG framework is able to account for the spatially varying flip angle across the brain, leading to $T_2$ maps with more homogeneous contrast (d). (a) simulated using an EPG framework (TE = 10 – 60 ms with a 10 ms echo spacing, $T_2$ = 30 ms), with resulting curves normalised to the signal at TE = 10 ms to aid visualisation of the deviation from a mono-exponential decay. Note that there is a degeneracy when $\alpha > 180°$, where flip angle $\alpha$ produces the same signal evolution as $360° - \alpha$.

$$\min_{B_1,T_2} \left\| \mathrm{TSE_{exp}}(x,y,z,\mathrm{TE}_{1:6}) - \mathrm{TSE_{sim}}(B_1,T_2,T_1,\mathrm{TE}_{1:6}) \right\|_2^2, \quad \text{[S1]}$$

where $\mathrm{TSE_{exp}}$ is the experimental TSE data over all six echoes ($\mathrm{TE}_{1:6}$) and $\mathrm{TSE_{sim}}$ is the simulated TSE signal using the EPG framework given a value of $B_1$, $T_2$, $T_1$ and the echo times. To avoid fitting for the signal amplitude ($S_0$), the experimental data and simulated signal were normalised (e.g. division by $\left(\sum_{j=1}^{6} \mathrm{TSE}_j^2\right)^{0.5}$). In a series of evaluations, $T_1$ was found to have very little effect on our $T_2$ estimates, after which it was set equal to a fixed

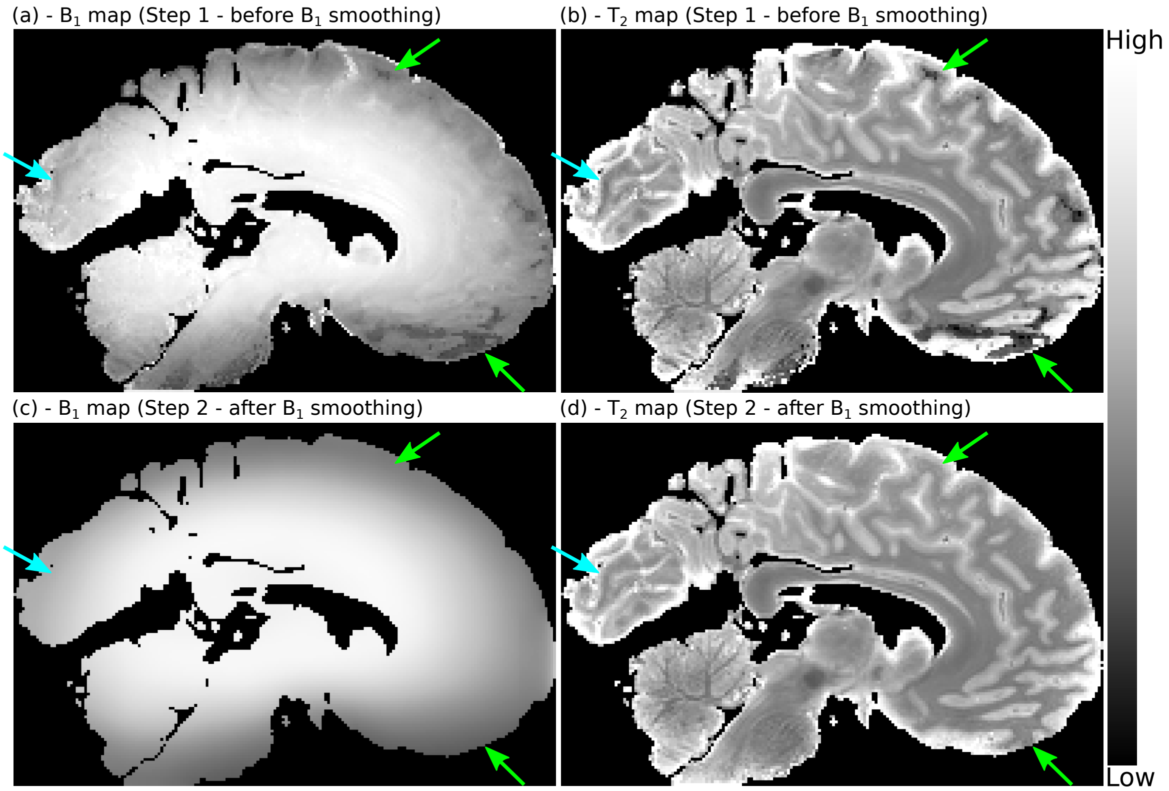


Figure S2: **$T_2$ estimates with EPG and smoothing $B_1$.** Step 1 - Our EPG model estimates $B_1$ maps a with decreasing $B_1$ as the brain boundary is approached (a), in addition to $T_2$ maps that do not have a strong dependence on the $B_1$ profile (b). The $B_1$ profile is expected to be smoothly varying across the brain. However, anatomical contrast is visible within the $B_1$ map (a – blue arrow), in addition to sharp discontinuities in areas close to the brain boundary (a - green arrows), regions associated with low SNR. This leads to artefacts and subtle contrast changes in the resulting $T_2$ map (b – blue and green arrows). Step 2 - By smoothing the $B_1$ maps with a polynomial filter (c) and fixing $B_1$ to the resulting map in a second stage estimate of $T_2$, we obtain consistent $T_2$ estimates across the brain (d). Here the $B_1$ maps (a and c) are displayed between 0 and 1, with the $T_2$ maps (b and d) displayed between 0 and 60 ms. N.B (d) does not include the regularisation step (Eq. [S3]).

constant (450 ms - the approximate value of measured $T_1$ in our postmortem datasets). Fitting was performed in MATLAB (version 2019b, The MathWorks, Inc., Natick, MA) based on code from (3) . Eq. [S1] was minimised using *lsqnonlin*.

Figures S2a and b show the resulting $B_1$ and $T_2$ maps for a single postmortem brain fit with this approach. The spatial inhomogeneity across the $T_2$ map is substantially reduced in comparison to the mono-exponential fit (Fig. S1c). However, $B_1$ maps are expected to vary smoothly across the brain and Fig. S2a reveals changes in $B_1$ that depend on the tissue contrast (e.g. blue arrow), in addition to underestimation of $B_1$ close to the brain boundary (e.g. green arrows). This leads to clear artefacts in the resulting $T_2$ maps (Fig. S2b), most apparent in regions where $B_1$ is underestimated (green arrows).

*Step two:*

To ensure spatial smoothness of $B_1$, the $B_1$ maps from step 1 were subsequently filtered using a local 3D polynomial filter (order = 2, kernel volume = 10x10x10 mm$^3$), with voxels weighted by the inverse of the standard error on the $B_1$ estimates. Figure S2c displays an example $B_1$ map after filtering, revealing a smoothly varying $B_1$ profile across the entire brain.

$T_2$ estimates were subsequently regenerated across the brain using the smoothed $B_1$ map as follows:

$$\min_{T_2} \left\| \mathrm{TSE}_{\mathrm{exp}}(x, y, z, \mathrm{TE}_{1:6}) - \mathrm{TSE}_{\mathrm{sim}}(B_1(x, y, z), T_2, T_1, \mathrm{TE}_{1:6}) \right\|_2^2, \qquad \text{[S2]}$$

where $B_1$ was fixed to the value of the smoothed $B_1$ map. Figure S2d displays the resulting $T_2$ map, where the most notable change is in regions of the brain where the $B_1$ was previously underestimated (Fig. 2d green arrows) – in these regions the $T_2$ estimates now match those of the surrounding tissue.

In regions very close to the brain boundary (characterised by very low $B_1$ and low SNR), in some brains the $T_2$ estimates were found to be highly sensitive to noise, leading to spurious values (Fig. S3a). To correct for this, regularisation was added to Eq. [S2] as follows:

$$\min_{T_2} \left\| \mathrm{W} \cdot [\mathrm{TSE}_{\mathrm{exp}}(x, y, z, \mathrm{TE}_{1:6}) - \mathrm{TSE}_{\mathrm{sim}}(B_1(x, y, z), T_2, T_1, \mathrm{TE}_{1:6})] \right\|_2^2 + \lambda \left\| \mathrm{T}_2 - T_{2,\mathrm{med}} \right\|_2^2, \qquad \text{[S3]}$$

where $\lambda$ is a regularisation constant ($\lambda = 2$), $T_{2,\mathrm{med}}$ is the median value of $T_2$ across the entire brain from the first step fitting and W is a scalar weight $\left(\sum_{j=1}^{6} \mathrm{TSE}_{\mathrm{exp,j}}^2\,(x, y, z)\right)^{0.5}$. Figure S3b displays a $T_2$ map with the addition of regularisation. Differences between the two maps (Fig. S3c) are restricted to these areas of very low $B_1$ and SNR. Figure 7 (Main Text) displays the $T_2$ maps fit using this approach for all 14 brains used in our study.

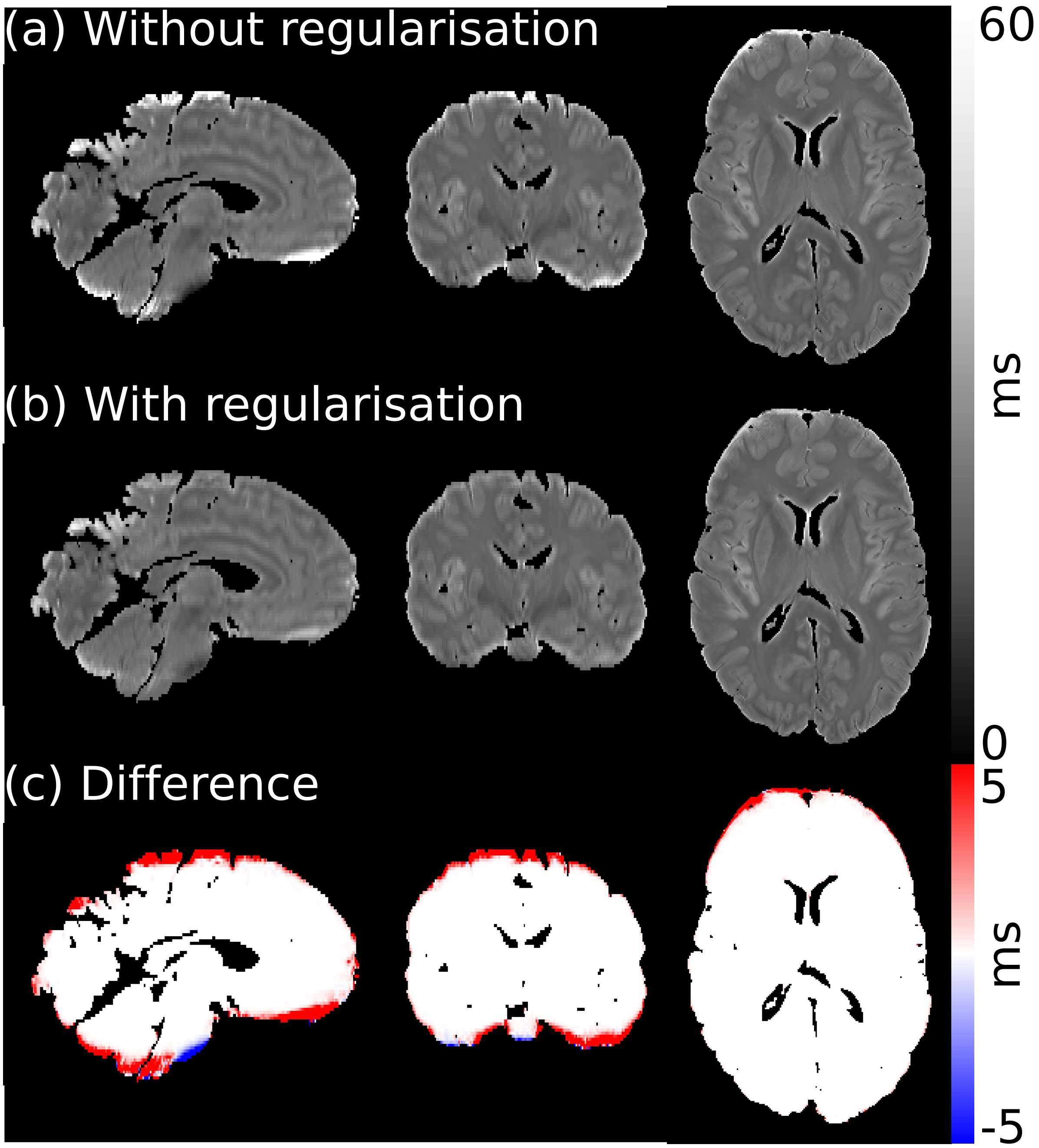


Figure S3: **Addition of regularisation for our $T_2$ estimates**. In areas of low $B_1$ in close proximity to the brain boundary, the TSE data had very low SNR. In some of our $T_2$ maps, this lead to spurious $T_2$ estimates (visible in (a)). The addition of regularisation (b) via fitting with Eq. [S3] bought the $T_2$ values within these regions into agreement with the surrounding tissue. Within other areas of tissue, the regularisation led to negligible changes in $T_2$ (c).

### Additional Supporting Figures and Tables

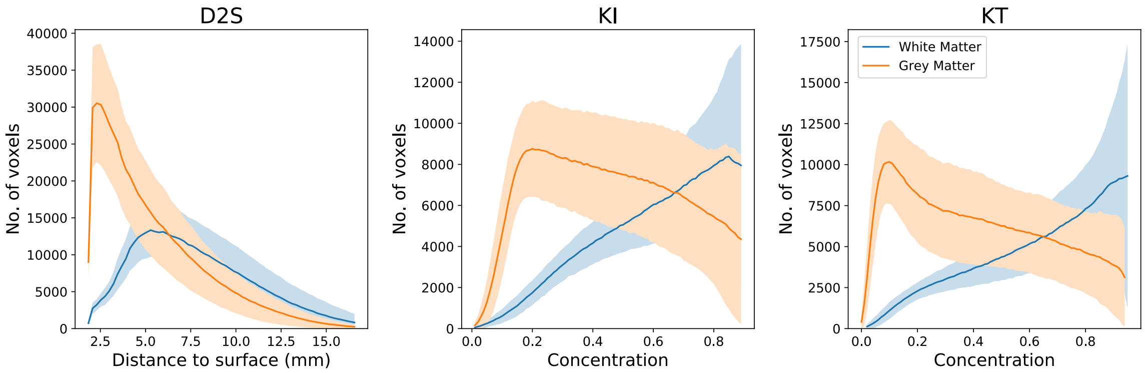


Figure S4: **Distribution of white / grey matter.** For all three models, white and grey matter are not evenly distributed with respect to distance from the nearest surface or fixative concentration. Plots generated over all 14 brains, with error bars representing the standard deviation between brains. Voxels < 2 mm from the nearest surface were not included in our analysis and are therefore not displayed here.

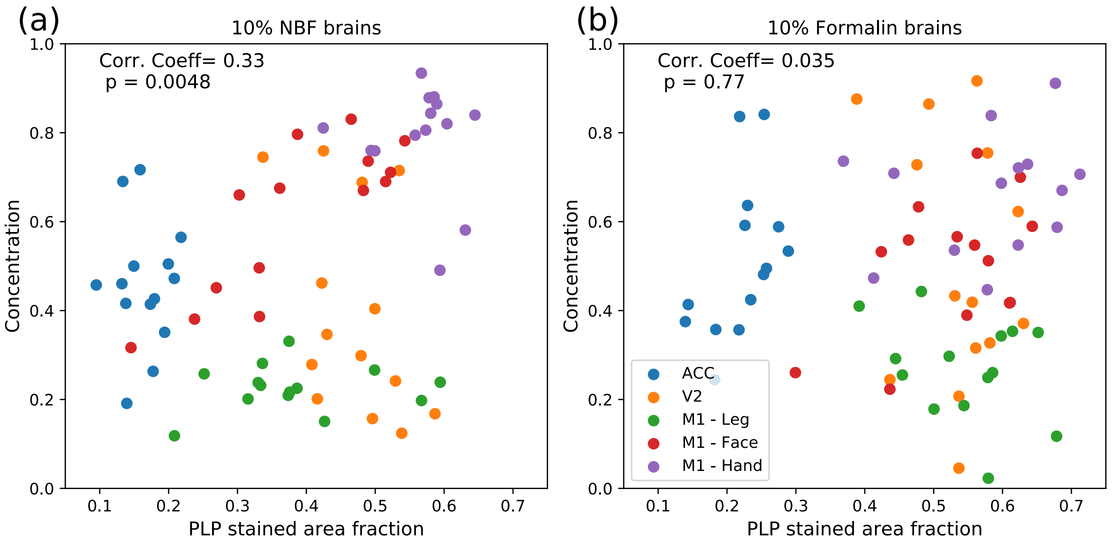


Figure S5: **Fixative concentration vs PLP SAF across M1, ACC and V2.** For brains fixed with 10% NBF, a small (but significant) correlation was found between the concentration of fixative and the PLP SAF, where there is a notable distinctions in the fixative concentration and PLP SAF in individual regions (e.g. ACC and M1 – hand region). For Brains fixed with 10% formalin, no such relationship was observed.

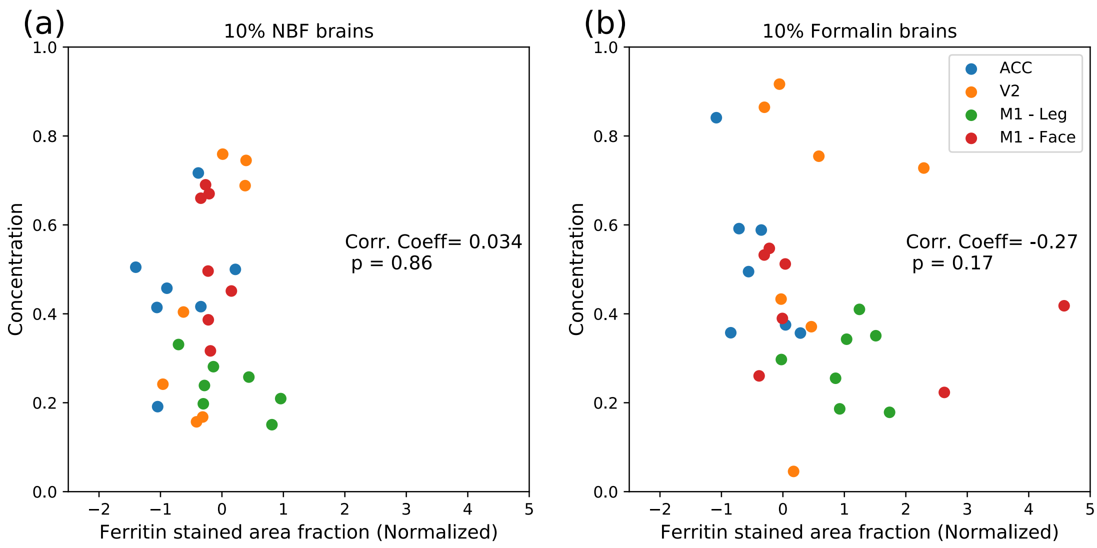


Figure S6: **Fixative concentration vs ferritin SAF across M1, ACC and V2.** No significant correlation was observed between fixative concentration and ferritin SAF for brains fixed with 10% NBF or 10% formalin. Note that as the ferritin SAFs were normalised for the two batches, the SAF values can be positive & negative and are not restricted to a range between 0 and 1. As the ACC and V2 regions were included in both batches, the ferritin SAFs were averaged prior to plotting.

(a) **10% NBF**

| *Tissue Type* | *Uncorrected* | *$B_1$ correction* |
|---|---|---|
| White Matter | $2.80 \pm 0.41$ | $2.56 \pm 0.34$ (0.26) |
| Grey Matter | $6.11 \pm 0.77$ | $5.79 \pm 0.70$ (0.43) |

(b) **10% Formalin**

| *Tissue Type* | *Uncorrected* | *$B_1$ correction* |
|---|---|---|
| White Matter | $1.46 \pm 0.23$ | $1.45 \pm 0.21$ (0.89) |
| Grey Matter | $2.45 \pm 0.32$ | $2.45 \pm 0.32$ (0.99) |

Table S3: **Inhomogeneity over white and grey matter for brains fixed with 10% NBF and 10% formalin – correction with $B_1$.** Correction with $B_1$ gave rise to a reduction in inhomogeneity across both grey and white matter for brains fixed with 10% NBF. However, this reduction in inhomogeneity did not reach significance, and was a smaller improvement vs the D2S, KI and KT models (Main Text Table 1). Little change was found in brains fixed with 10% formalin, consistent with the corrections using the D2S, KI and KT models (Main Text Table 1).

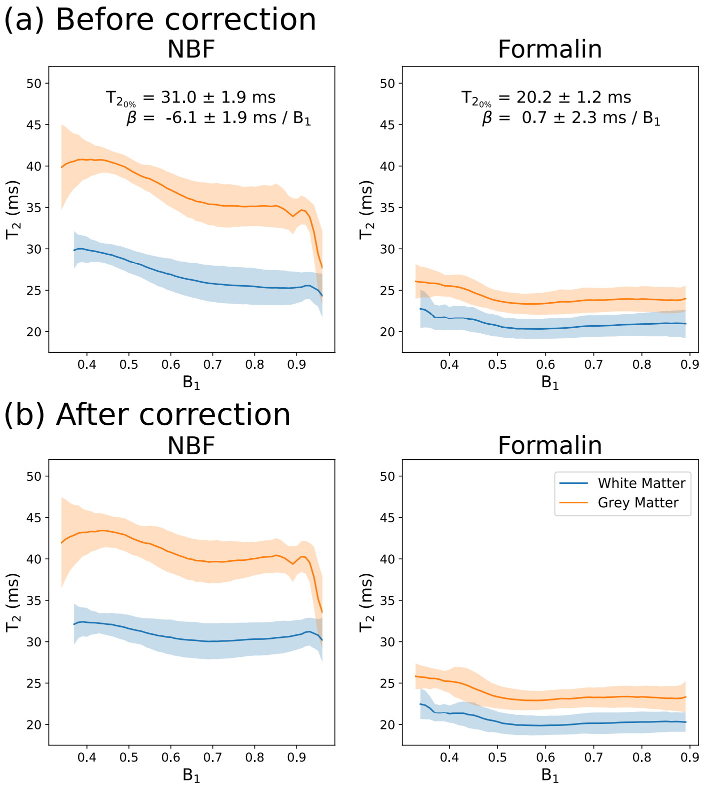


Figure S7: **$T_2$ vs $B_1$ over white and grey matter for all post-mortem brains fixed with 10% NBF and 10% formalin.** The corrections presented in this work assume that presence of fixative in brains is the source of variation in $T_2$ across the brains; however, the predicted patterns of fixative are similar to the spatial pattern of $B_1$. Here, we consider how much of the variation in $T_2$ may be explained by the $B_1$ spatial profile. A dependency is observed vs $B_1$ in our post-mortem cohort (a), with a stronger effect in brains fixed with 10% NBF (left) vs 10% formalin (right), consistent with the other three models investigated in this study (Main Text Figs. 9 and 10). However, regressing out the influence of $B_1$ using Eq. [6] (b) leads to a higher remaining inhomogeneity as shown in Table S3 vs the D2S, KI and KT models for the brains fixed with 10% NBF (Main Text Table 1) and similar performance (no change) for brains fixed with 10% formalin. Results displayed as the mean ± standard deviation across brains.
